# Tropicalisation of rocky shore gastropod communities along the Baja California Peninsula

**DOI:** 10.1101/2025.11.20.689443

**Authors:** Karolina M. Zarzyczny, David A. Paz-García, Suzanne T. Williams, Marc Rius, Moira MacLean, Phillip B. Fenberg

## Abstract

Tropicalisation is rapidly restructuring marine communities globally. Despite extensive research in subtidal habitats, the extent of tropicalisation and its impact on rocky intertidal communities remains poorly understood. Here we integrated contemporary field surveys with genetic barcoding, historic surveys, museum collections, and fossil records to document changes in the rocky shore gastropod communities along 8 degrees of latitude (approximately 3,120 km of coastline) of the Baja California Peninsula. We detected poleward range expansions of 30 tropical and subtropical species, and trailing-edge retractions of 13 temperate species, with some range shifts occurring across long-standing biogeographic boundaries. Fossil evidence reveals that several range-extending species previously occupied latitudes beyond their modern distributions during Pleistocene warm periods, suggesting that contemporary expansions echo historical responses to climatic warming, albeit under accelerated rates. Using *Lottia gigantea* as a case study, we demonstrated early range retractions can be marked by shifts in size-frequency distributions demonstrating declines in abundance, and loss of small individuals suggestive of reduced recruitment. We showed that rocky intertidal systems are sensitive to climate-driven range shifts, and highlighted the importance of integrating historical, palaeontological and contemporary data to detect early ecological and evolutionary consequences of tropicalisation. Continued monitoring combined with molecular approaches is crucial for predicting and managing biodiversity responses under ongoing climate change.

## Introduction

Climate-inducted range shifts are rapidly restructuring global biodiversity patterns as the spatial distribution of species shift to track suitable climatic conditions (Fredston-Hermann et al., 2020; Jacobsen, 2020; Ramalho et al., 2023; Rubenstein et al., 2023; Zarzyczny et al., 2024a). For marine ecosystems, climate-induced range shifts are especially prominent as marine species tend to better track climate niches than terrestrial species (Burrows et al., 2019; Colwell & Feeley, 2025; Lenoir et al., 2020; Pinsky et al., 2020). At mid- to low-latitudes in particular, many tropical species are expanding their ranges poleward, whilst temperate species are experiencing range retractions at the trailing-edge of their distributions (Zarzyczny et al., 2024a). This globally documented phenomenon has been coined ‘tropicalisation’ (Vergés et al., 2014; Wernberg et al., 2013; Zarzyczny et al., 2024a).

Tropicalisation has the potential to fundamentally reshape community structure, ecosystem functioning and services provided by marine ecosystems (Zarzyczny et al., 2024a). Shifts in species interactions, such as altered grazing pressure (Kumagai et al., 2018; Vergés et al., 2016; Zarzyczny et al., 2022a) or changes in predation (Fenberg et al., 2023) can have profound ecological and evolutionary consequences (Zarzyczny et al., 2024a). Functionally, tropicalisation can lead to changes in primary production (Peleg et al., 2020; Vergés et al., 2014), nutrient cycling (Macy et al., 2021; Spatharis et al., 2012; Sullivan et al., 2021), and habitat complexity, particularly where habitat-forming species are affected (Vergés et al., 2014). Beyond ecological impacts, tropicalisation can also have cascading socioeconomic consequences, altering the availability of fisheries resources (Cheung et al., 2013; Yamamoto et al., 2020), influencing tourism and coastal livelihoods (Sudo et al., 2022). Documenting and understanding tropicalisation is therefore essential for predicting and managing the future of marine ecosystems under climate change.

To date, tropicalisation research has focussed predominately on subtidal habitats, particularly on shallow reefs including macroalgal beds or coral communities (Vergés et al., 2016; Wernberg et al., 2013). In contrast, rocky intertidal systems have received comparatively little attention despite their ecological importance and vulnerability (Hawkins et al., 2009; Mieszkowska et al., 2021). Intertidal rocky shores are particularly vulnerable to climate change due to their exposure to both marine and aerial thermal stress, ultraviolet radiation and other environmental changes (Marshall et al., 2015; Przeslawski et al., 2005; Stafford et al., 2015). Moreover, the fragmented nature of many rocky shore habitats makes resident species particularly vulnerable to climate impacts, as opportunities for range shifts may be constrained by patchy availability of suitable substrata and/or the species’ dispersal capabilities (Zarzyczny et al., 2025; Zarzyczny et al., 2024b). Long-term studies have demonstrated that rocky shores are highly responsive to climate-driven change, with shifts in species distributions and community structure documented in temperate systems (Hawkins et al., 2009; Mieszkowska et al., 2021; Sanford et al., 2019). However, the role of tropicalisation in shaping rocky shore assemblages at tropical-temperate biogeographic transition zones, remains largely unexplored (but see Zarzyczny et al., 2024b).

Whilst tropicalisation refers to contemporary shifts in species distribution (Zarzyczny et al., 2024a), similar distributional changes have occurred in the deeper past (Hellberg et al., 2001; Hurtado et al., 2007). Pleistocene (1.8 million to 10,000 years ago) climatic oscillations, characterised by alternating interglacial warm phases and glacial maxima, provide a case study of past population dynamics in response to climate change (Hofreiter & Stewart, 2009). The repeated range expansions and contractions have left a complex but lasting imprint on the genetic structure and biogeography of extant taxa (Hellberg et al., 2001; Hofreiter & Stewart, 2009). Notably, some taxa, such as the marine gastropods *Nerita funiculata* and *N. scabricosta*, exhibit genetic signatures of extensive range expansions during the Pleistocene (Hurtado et al., 2007), and are once again undergoing poleward range expansions linked to the tropicalisation of the Baja California Peninsula (Zarzyczny et al., 2024b). This recurring trend suggests that present-day tropicalisation of rocky shores may echo past responses to climatic warming, albeit under the accelerated and sustained trajectory of contemporary climate change. However, it remains unknown whether other currently range-expanding species exhibit a similar pattern of repeated expansion dynamics observed during Pleistocene warm periods.

Among rocky shore gastropods, the temperate limpet, *Lottia gigantea*, provides a particularly informative case study for understanding climate-driven range dynamics on the Eastern Pacific (EP) coast, and the ecological implications of tropicalisation. *Lottia gigantea* is a large, grazing limpet distributed along the EP coastline from Baja California, north to northern California (Nielsen et al., 2024; Sanford et al., 2019). It is the largest limpet species found in North America, with shell length reaching 100 mm in the largest individuals (Kido & Murray, 2003). Given their large size and territorial grazing behaviour, *L. gigantea* are ecosystem engineers as they prevent recruitment of other intertidal species on their occupied substrate (Kido & Murray, 2003; Stimson, 1970). However, the large size also makes *L. gigantea* particularly attractive to human harvesting (Fenberg & Roy, 2012). As a result, *L. gigantea* has been a subject to more scientific investigation than many other intertidal gastropods, providing us with valuable baseline data for ecological study. Furthermore, evidence suggests that *L. gigantea* populations are already affected by climate change. In the northern portion of this species’ range (northern California), *L. gigantea* has experienced increased recruitment associated with the 2014-2016 marine heatwaves (Sanford et al., 2019), potentially signalling early stages of poleward range expansion under warming conditions. Meanwhile, southernmost populations of *L. gigantea* are characterised by unique genetic diversity (Nielsen et al., 2024) which could be lost if the species was to undergo a southern range retraction. Taken together, *L. gigantea* provides a valuable model for detecting early signs of tropicalisation.

Here, we first combine historical records from museum collections, online data repositories, and past surveys of 73 gastropod species, with contemporary field surveys to document tropicalisation of rocky shore gastropod communities on the Baja California Peninsula coast over the last 24 years. To date, only a handful of species range shifts have been documented on rocky shores of the Baja California Peninsula (Fenberg et al., 2014, 2023; Zarzyczny et al., 2024b). Secondly, we utilise fossil records to determine whether range expanding species show recurring patterns of poleward expansion associated with Pleistocene warming. Finally, we examine size-frequency distributions of one of the best studied species, *L. gigantea*, at the trailing-edge of its distribution comparing recent surveys with those conducted approximately 20 years earlier to investigate changes in population demography and range shifts.

## Methodology

### Study region

The Baja California peninsula in Northwestern Mexico is bordered by the Pacific Ocean to the west (i.e. the eastern Pacific (EP) coast), and the Gulf of California (GoC) to the east (Figure 1). The peninsula stretches across three distinct marine biogeographic regions on the EP coast; the temperate Southern California Bight, the Magdalena Transition Zone, and the Mexican Tropical Pacific, which respectively host a range of cool and temperate (including warm temperate), subtropical, and tropical species (Fenberg & Rivadeneira, 2019; Lluch-Belda et al., 2003; Zarzyczny et al., 2024b). On the east coast of the peninsula, the GoC predominantly hosts subtropical and tropical species, with some occurrences of warm temperate species in the head of the Gulf (Aceves-Medina et al., 2004; Fenberg & Rivadeneira, 2019; Zarzyczny et al., 2024b). Meanwhile, the alternating rocky and sandy shores of the Baja California Peninsula further shape the coastal biogeographic structure (Fenberg et al., 2015; Fenberg & Rivadeneira, 2019).

**Figure 1.**
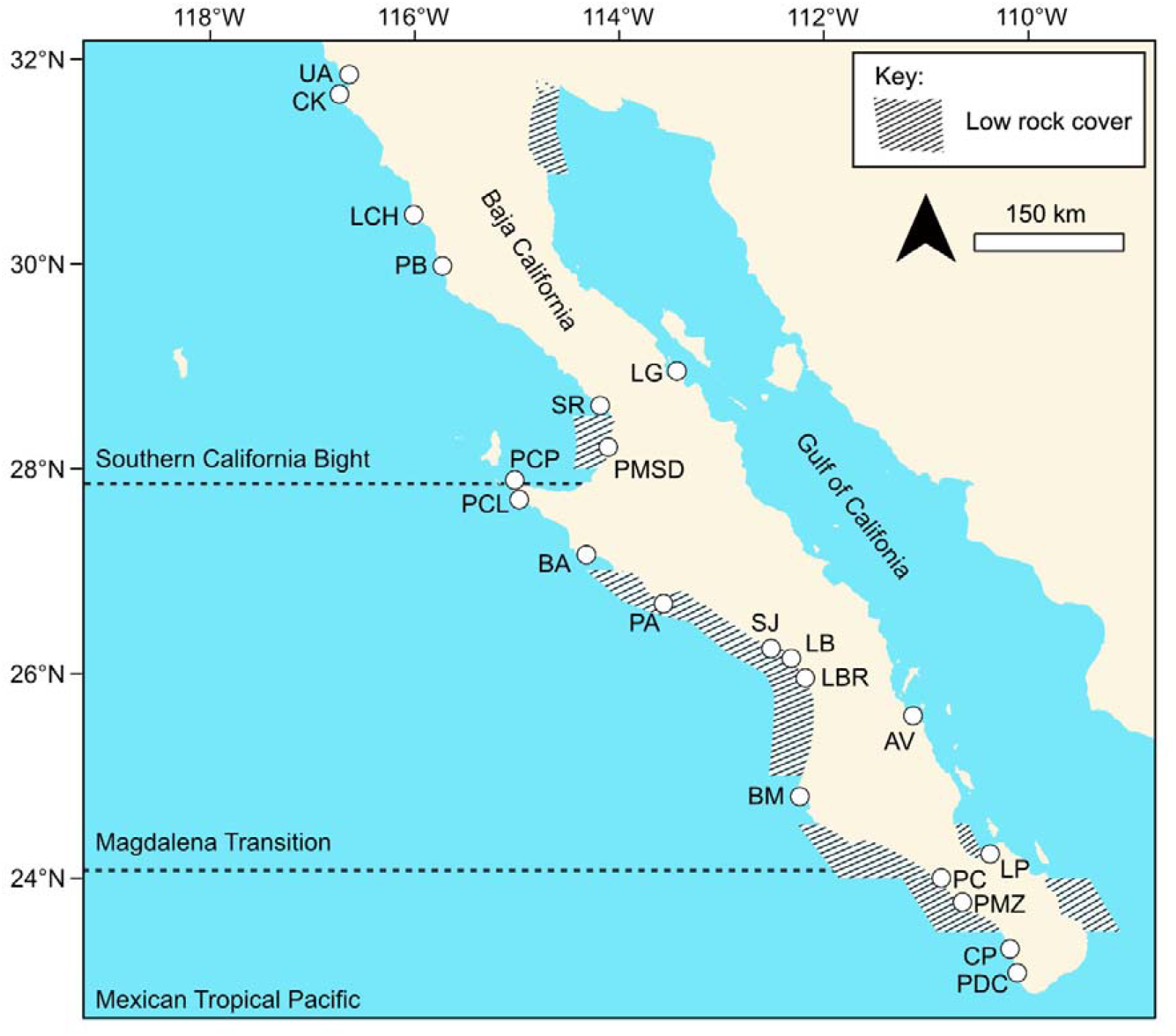
Map of Baja California Peninsula demonstrating the marine biogeographic regions adapted from (Blanchette et al., 2008; Fenberg et al., 2015; Spalding et al., 2007); Low rock cover data for the peninsula only, was extracted from (Fenberg & Rivadeneira, 2019), and is defined as <20% rock cover per 0.5°. The 18 study sites are LG = La Gringa, AV = Agua Verde, LP = La Paz, PDC = Pozo de Cota, CP = Cerritos Point, PMZ = Punta Marquez, PC = Punta Conejo, BM = Bahía Magdalena, LBR = Las Barrancas, LB = La Bocana, SJ = San Juanico, PA = Punta Abreojos, BA = Bahía Asunción, PCL = Punta Clambey, PCP = Punta Caballo de Piedra, PMSD = Punta Morro Santo Domingo, SR = Santa Rosalillita, PB = Punta Baja, LCH = La Chorera, CK = Campo Kennedy, and UA = Universidad Autónoma de Baja California (UABC).

Unlike in many tropicalisation hotspots where persistent, poleward-flowing west-boundary currents such as Kuroshio current (Japan) or Eastern Australia current, facilitate tropicalisation (Kumagai et al., 2018; Messer et al., 2020; Vergés et al., 2014; Zarzyczny et al., 2022b), Baja California is largely influenced by the eastern-boundary California Current (Checkley & Barth, 2009). In winter and spring, this cool-southward-flowing current promotes upwelling and maintains temperate conditions (Durazo, 2015). During summer and autumn, the region is influenced by an inflow of tropical and subtropical waters, and the intensification of the poleward-flowing counter current (Checkley & Barth, 2009; Durazo, 2015), which could support tropicalisation. Furthermore, during El Niño events warm anomalous poleward currents form along the coast facilitate tropicalisation of the peninsula (La Rosa-Izquierdo et al., 2022). These seasonal and episodic intrusions of warm water, contrast with the persistent flow seen western boundary systems, creating more transient and variable opportunities for tropicalisation along the Baja California Coast.

### Establishing species ranges

To document tropicalisation, we surveyed 18 sites along the EP coast, and three sites within the GoC (Figure 1; Table S1) between September and October 2017. This initial survey was followed by more in March 2018, December 2021, January 2022, and January 2024. We carried out exhaustive surveys at low tide, identifying all encountered gastropod species, but restricting each survey to ∼2h. Species that we were unable to identify in the field were photographed, and immediately preserved in 70% ethanol (EtOH), before being preserved in molecular grade EtOH in the laboratory, until DNA extraction. Gastropods with an operculum were gently cracked, or had their operculum pierced before being placed in EtOH to ensure rapid preservation of the tissue. Using the survey data, we estimated the northern range limits for tropical and subtropical species, and southern range limits for temperate species. Where possible, we refined the location of the range limits using independent species entries on iNaturalist (https://www.inaturalist.org). In such cases, geographic coordinates were extracted for inclusion in our database after species identification was confirmed for each entry using associated photographs of the records.

For all encountered gastropod species, we determined the previously recorded historic range limits (recorded in the year 2000 or earlier, unless stated otherwise). We consulted primary literature, books, online repositories such as Global Biodiversity Information Facility (GBIF, www.GBIF.org), historic field surveys, accessed museum collections and consulted museum curators. Where available, we extracted fossil records of our species of interest from GBIF (www.GBIF.org), to estimate the maximum range limits of tropical species during the Pleistocene (2.6 million to 11,700 years ago) climatic fluctuations (Hofreiter & Stewart, 2009). Undated fossil records, records outside of the study area, and those with no locality information (e.g. where country of collection was the only geographic information provided) were excluded from the study.

To estimate the extent of modern range changes, we plotted the historic and modern range limits of all detected gastropod species, onto the coastal transect produced by Fenberg & Rivadeneira, (2019). The transect fragment utilised in this study ranges from the head of the GoC, south to the tip of the Baja California Peninsula, and north along the EP coastline (Figure 1). Briefly, the transect was generated in cumulative 48 km bins, by tracing the contour of the coastline from 5 km above the sea level using Google Earth (v 7.1.5; Fenberg & Rivadeneira, 2019). Where the range limit occurred in between two 48 km bins, the mid-point between the two bins was utilised for the calculation of a range change. Where a range expansion occurred from the GoC to the EP coastline, the estimated distance of range expansion was calculated from El Cardoncito (23.22° N, 109.45°W) at the tip of the GoC. In the case of La Bocana and San Juanico which fall within the same 48 km bin, we followed the original methodology described by the authors to estimate the coastal transect between the two sites.

### DNA barcoding for cryptic species diversity

For species which we could not identify based solely on morphology, or those which harbour cryptic diversity (i.e. distinct species which have historically been classified as a single species due to morphological similarities), we extracted DNA from foot tissue with the DNeasy Blood and Tissue Kit (Qiagen) following a modified protocol described by Zarzyczny, Hellberg, et al., (2024). For species with mucus-rich tissue and high polyphenolic content that can clog spin columns (e.g. Littorinids), we used the CTAB extraction method described by Williams et al., (2003). For all species, we amplified a fragment of Cytochrome Oxidase Subunit I (COI) using primers, reactions and optimal PCR conditions outlined by (Zarzyczny et al., 2024b). We purified PCR products using the QIAquick PCR Purification Kit (Qiagen), according to manufacturer instructions, and sequenced them using Sanger Sequencing (Eurofins Genomics). We used the sequences to confirm species identification by comparing partial COI barcodes among morphotypes (Hollister et al., 2023), and with GenBank entries (https://www.ncbi.nlm.nih.gov/genbank/) using Nucleotide BLAST (https://blast.ncbi.nlm.nih.gov/Blast.cgi).

### Lottia gigantea *case study*

We compared abundance and size data for *Lottia gigantea* from surveys conducted in 2003-2006 (Fenberg, 2008) with new data from 2021-2024, which was collected using the same methodology. We carried out surveys at the four sites comprising the species’ southern range limit (La Bocana, San Juanico, and Punta Abreojos). At each site, we conducted a two-hour search during low tide and measured all *L. gigantea* individuals encountered during the two-hour period. Shell-length, defined as the maximum distance from the anterior to the posterior end of the shell, was measured using callipers to the nearest millimetre.

We conducted a statistical analysis to test whether the average limpet size changed between 2003 and 2024, at each site. As the data violated the assumption of normality, we conducted a Mann-Whitney U test with p-values adjusted for multiple comparisons using the Bonferroni method. All data analysis was conducted using RStudio version 4.5.1.

## Results

### Field surveys

We detected a total of 99 species of gastropods across the rocky coastline of the Baja California Peninsula (Table S2). We describe in more detail the distribution of all encountered species in ‘Species Accounts’ in Supplementary Information S1. We encountered occasional records of gastropods which we were unable to confidently identify to a species level or obtain a reliable COI barcode for identification, and subsequently we excluded these from further study. These records included *Costoanachis* sp. (La Gringa), *Eulithidium* sp. (La Bocana and Punta Abreojos), *Fissurella* sp. (Punta Marquez), *Mitrella* sp.1 (La Gringa), and Vermetidae sp. (San Juanico). Furthermore, using the partial COI marker gene, we were unable to genetically distinguish five morphologically similar *Echinolittorina* species, at the species level. Although the species separated into two distinct genetic clusters, species boundaries within each cluster remained unclear due to high genetic similarity. The first cluster includes *E. aspera, E. dubiosa*, and *E. tenuistriata*, while the second cluster includes of *E. apicina* and *E. paytensis*. As species delimitation falls beyond the scope of this study, from herein we refer to the two clusters as *E. aspera* group and *E. apicina* group, respectively (Supplementary Information S1).

We detected occasional records of several gastropod species for which we were unable to determine the modern distribution. For example, during our rocky intertidal surveys, we observed several tropical and subtropical species including *Columbella aureomexicana, C. fuscata, Conus nux, Coralliophila nux, Eupleura muriciformis, Heliacus areola, Neorapana tuberculata, Neoterebra variegata, Polinices uber, Rissoina stricta, Zetecopsis zeteki*, all of which also occupy the subtidal region (Supplementary Information S1). Similarly, we detected occasional records of species which typically occupy habitats other than rocky shores such as *Macron aethiops, Modulus cerodes, Notocochlis chemnitzii, Parvanachis diminuta*, which demonstrate a preference for mudflats, or *Vitta luteofasciata* which frequently inhabits mudflats and mangrove swamps. We also detected several temperate species such as *Callianax biplicata, Norrisia norrisii, Pomaulax gibberosus*, and *Trachypollia lugubris*, which have distributions extending into the subtidal region (Supplementary Information S1). As the distribution of these species extends beyond the scope of our surveys, we are unable to reliably capture their complete modern distribution, and consequently we have excluded these species from our study.

### New geographic range boundaries

After excluding uncertain records, we report modern distributions (and where possible historic distributions) for 49 subtropical and tropical species (Table 1), and 24 temperate species of rocky shore gastropods (Table 2). We detected poleward range expansions for 30 subtropical and tropical species (Table 1; Figure 2), and 13 trailing-edge range retractions of temperate species (Table 2; Figure 3). For the 30 subtropical and tropical species, the range expansions ranged from ∼48 km to ∼1008 km, with an average range expansion of ∼403 km. For the 13 temperate species, the range retractions ranged from ∼33 km to ∼648 km, with an average range retraction of ∼237 km.

**Table 1.**
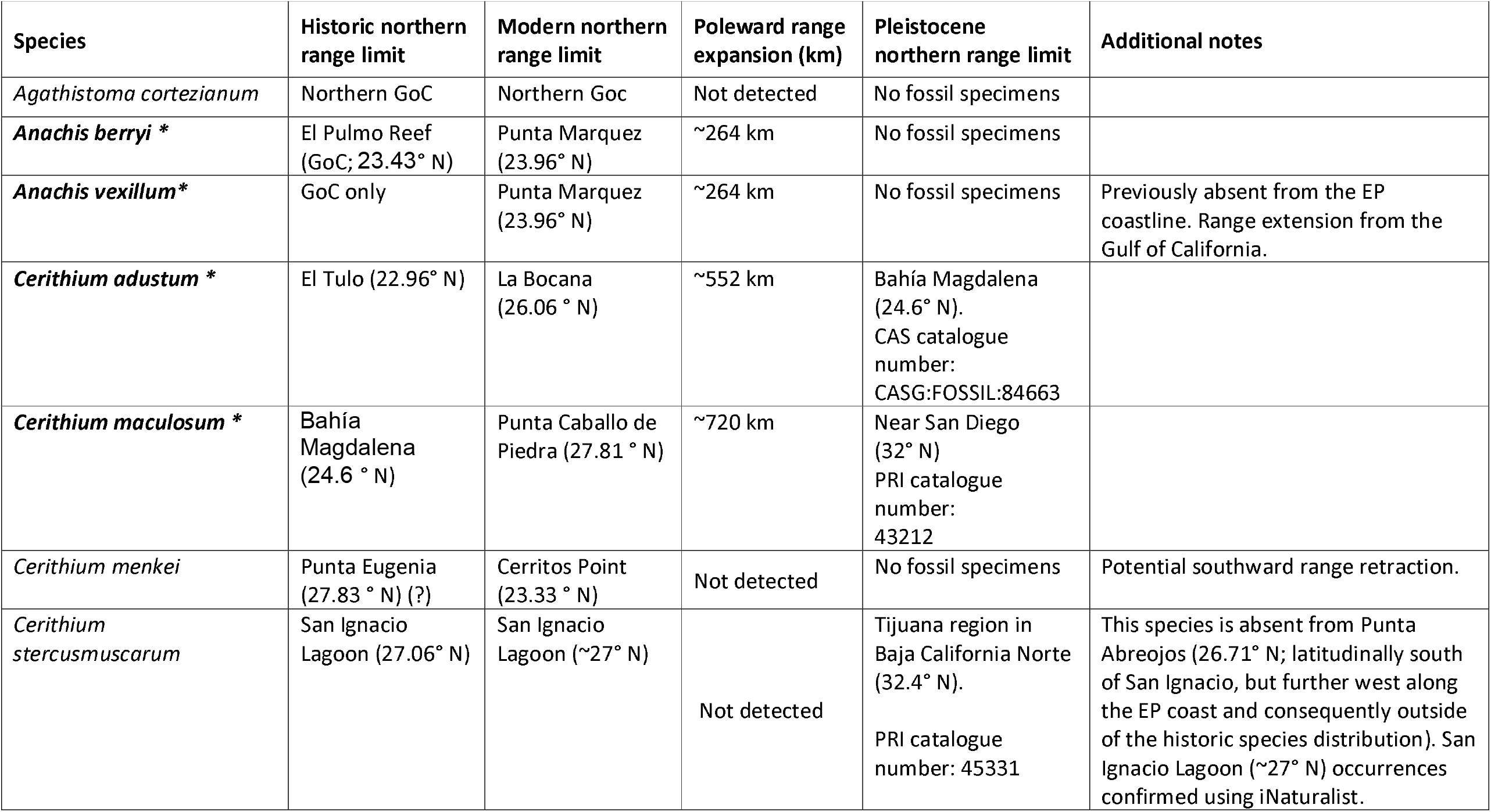
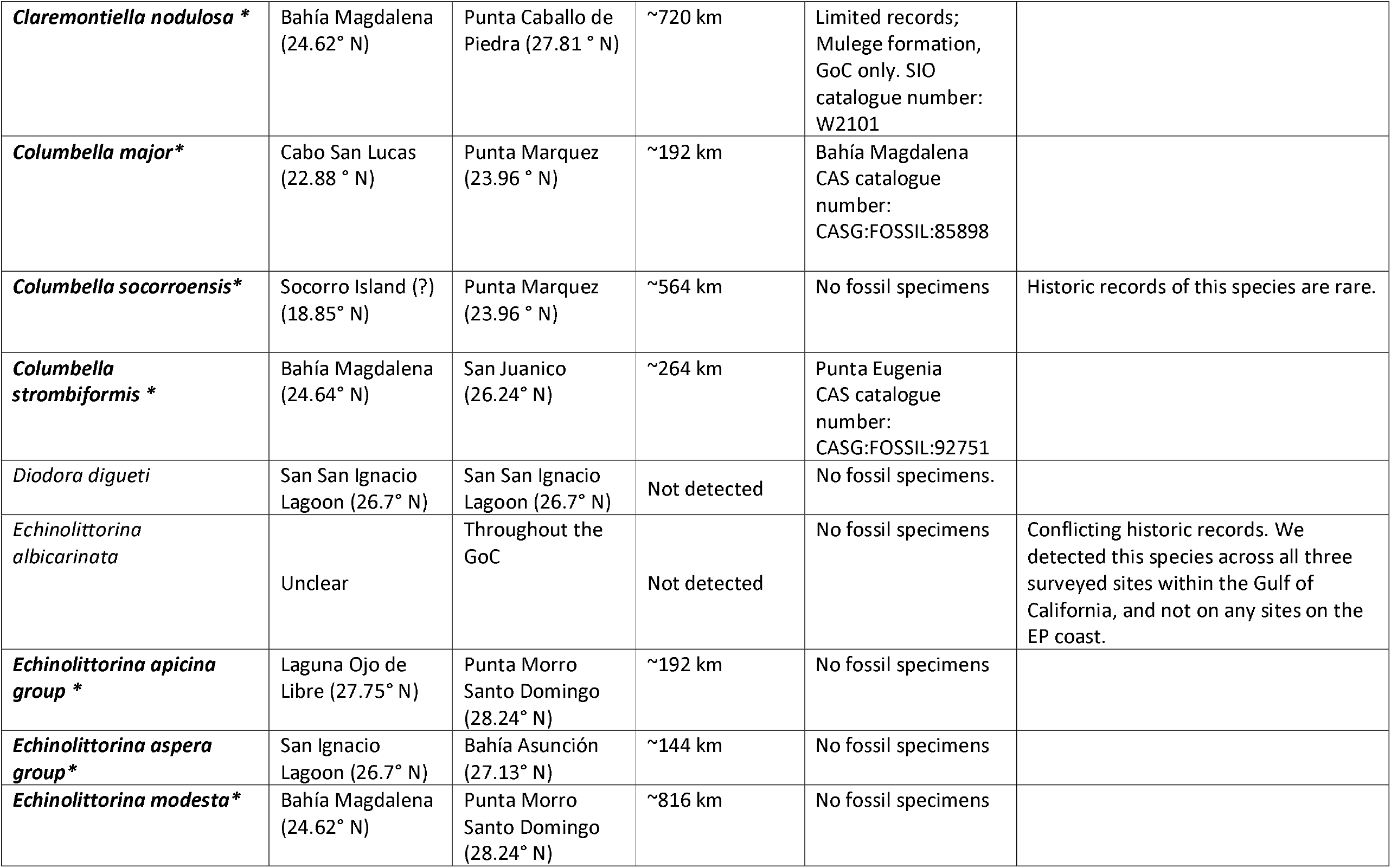
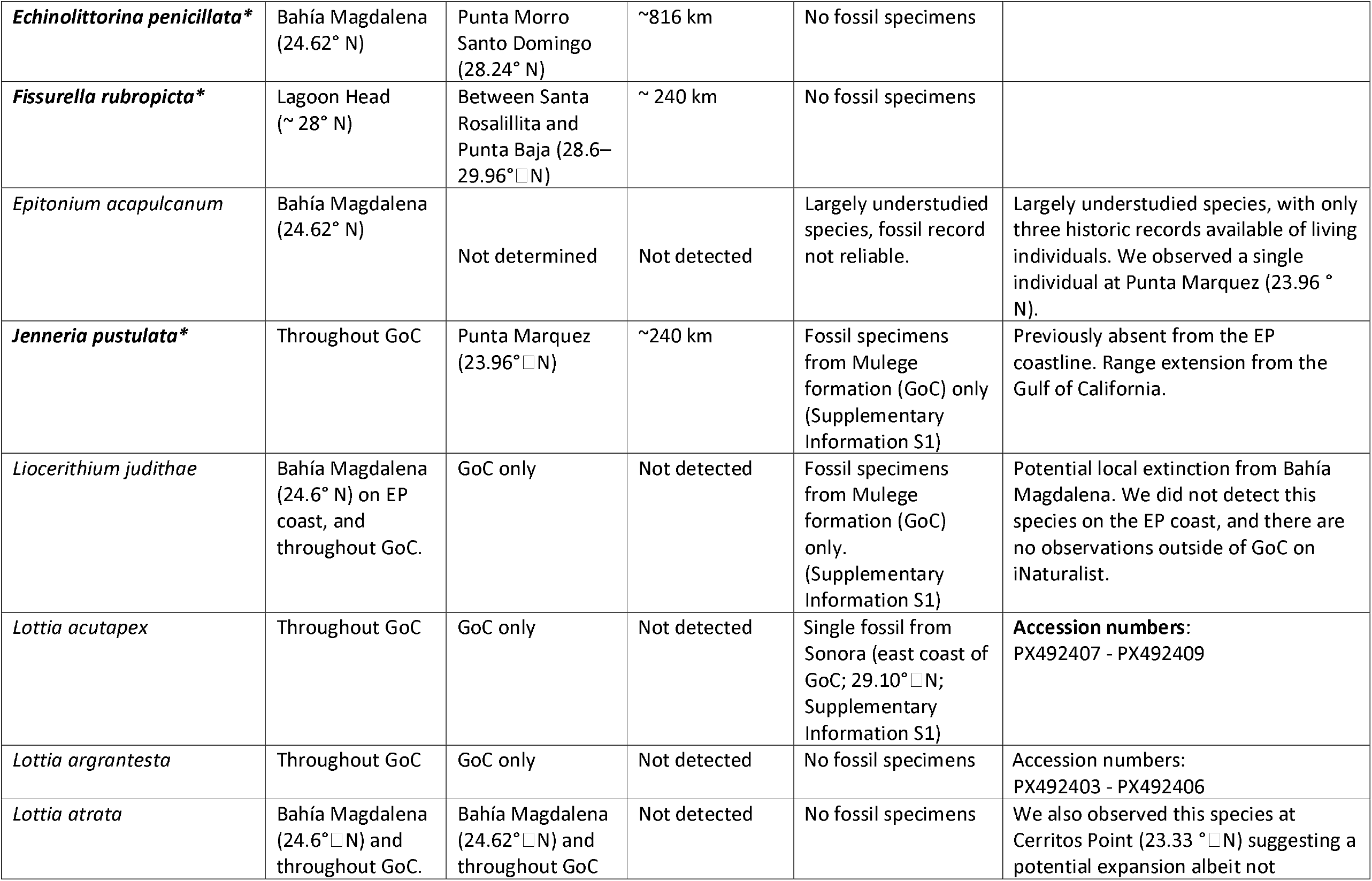
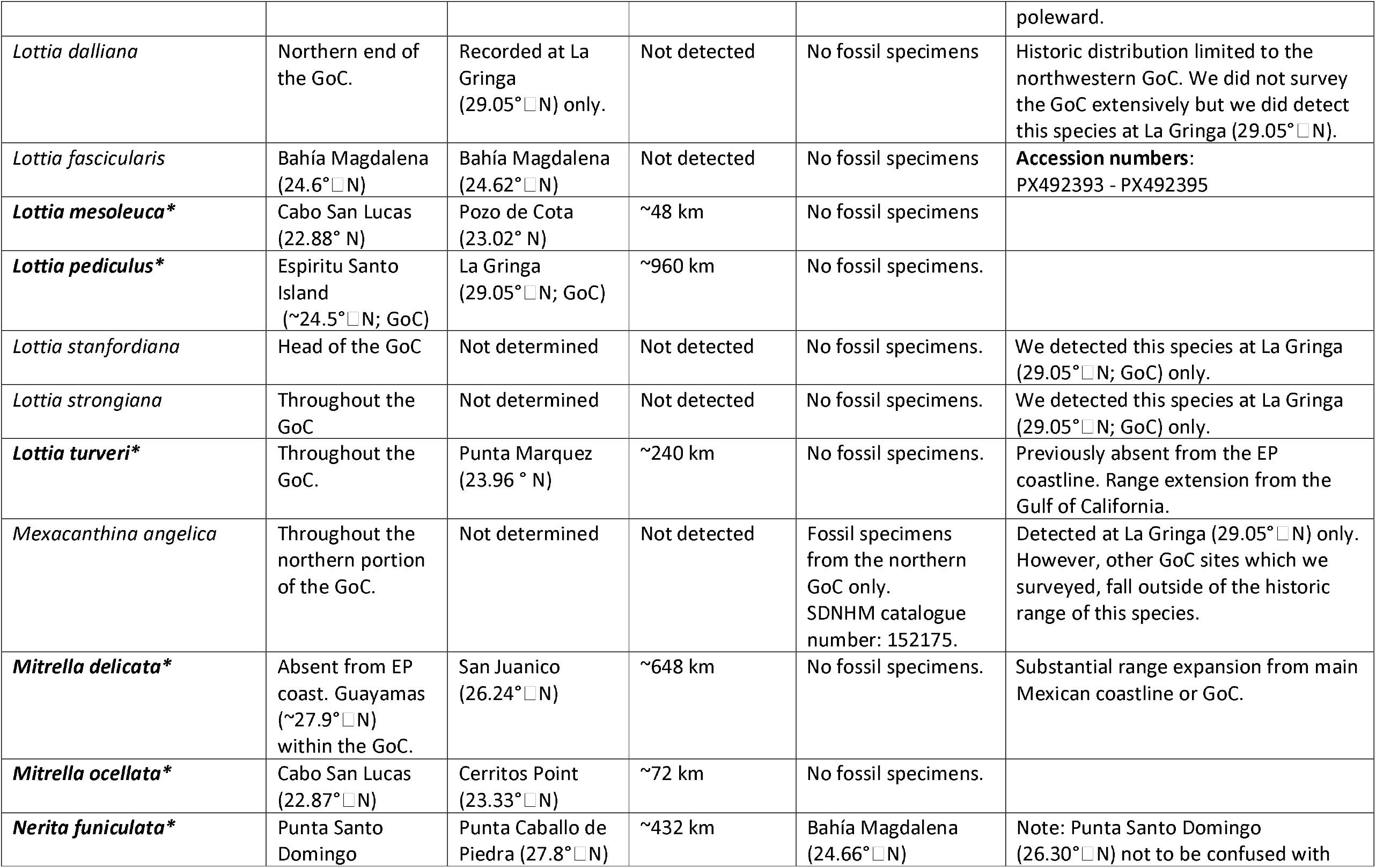
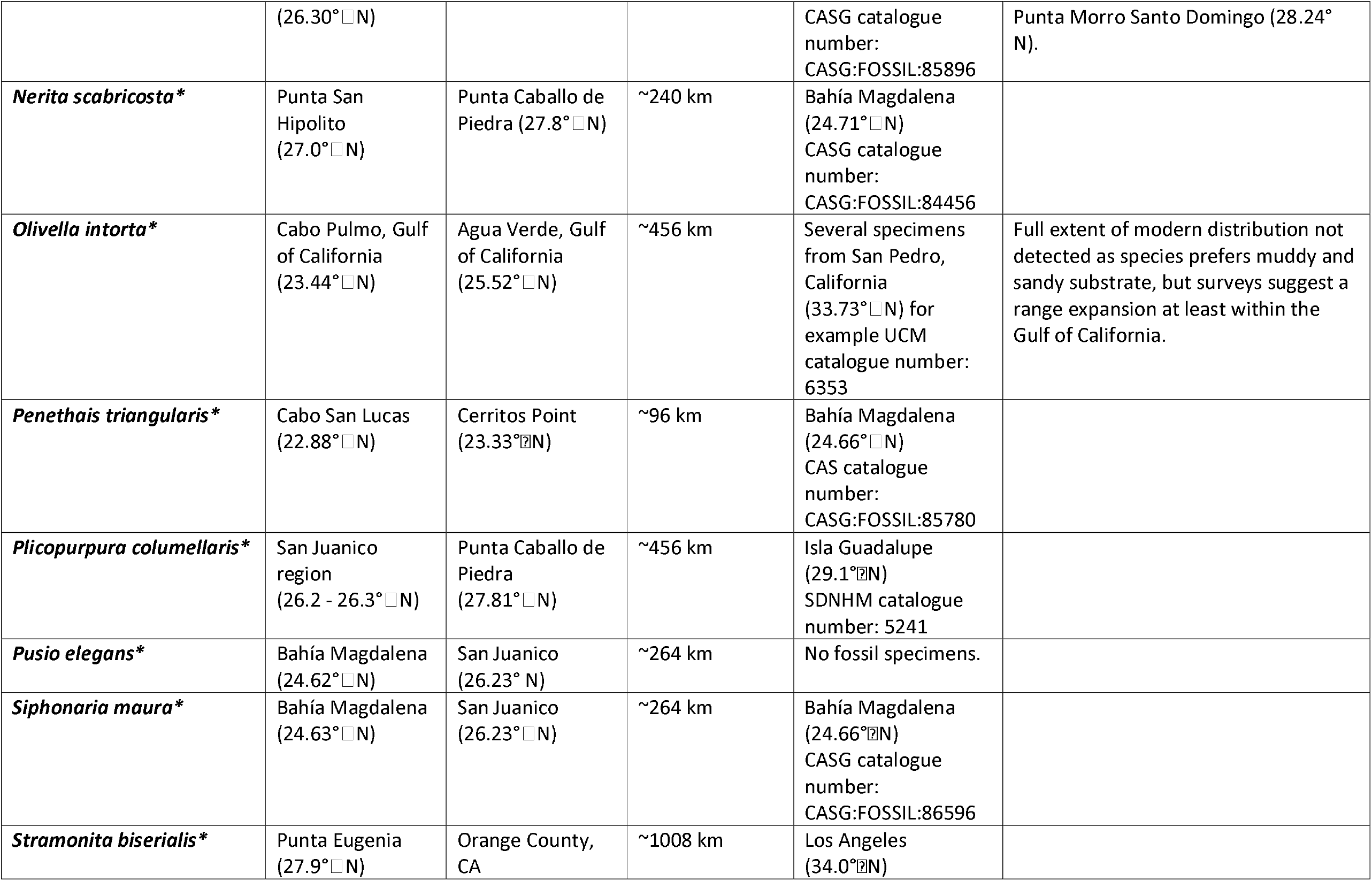
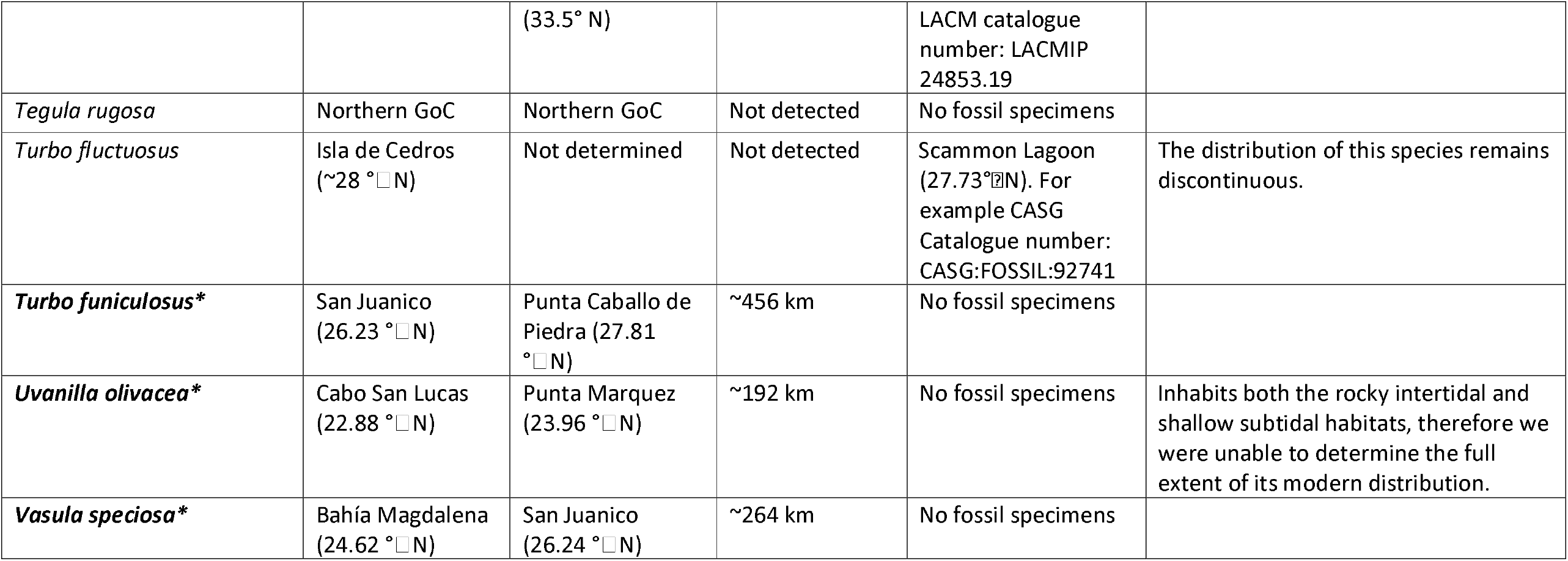
Modern and historical northern range limits and range shift in km of tropical rocky shore invertebrates. Range expanding species are bolded and marked with *. Detailed description of species ranges, and relevant historic species records are described in Supplementary Information S1. Where plicable we provide the Accession Numbers to the partial COI genetic markers obtained in the study). (?) = indicates uncertain historic range limit. GoC = Gulf of California; CAS = California Academy of Sciences; PRI = Palaeontological Research Institution.

**Table 2.**
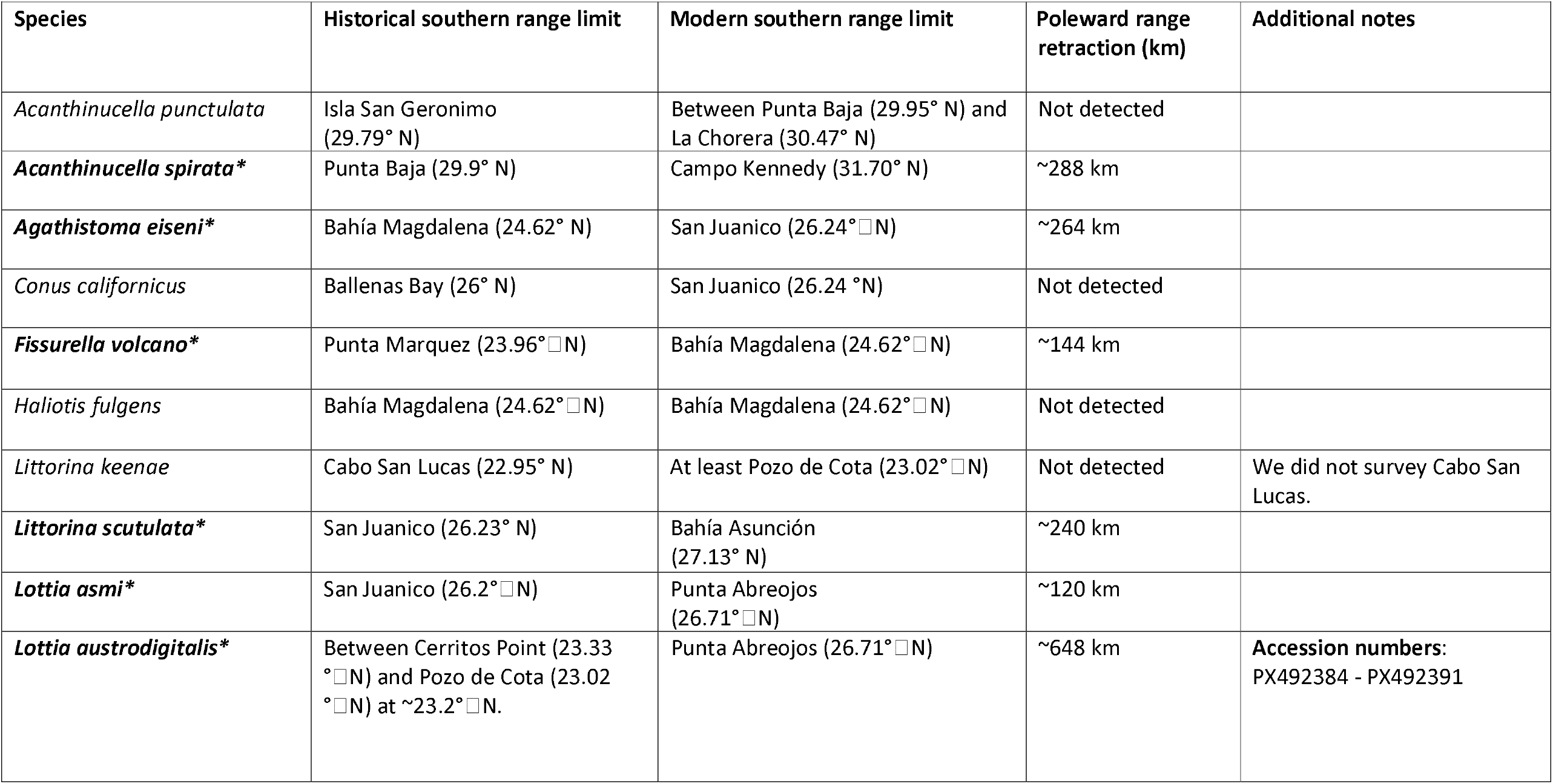
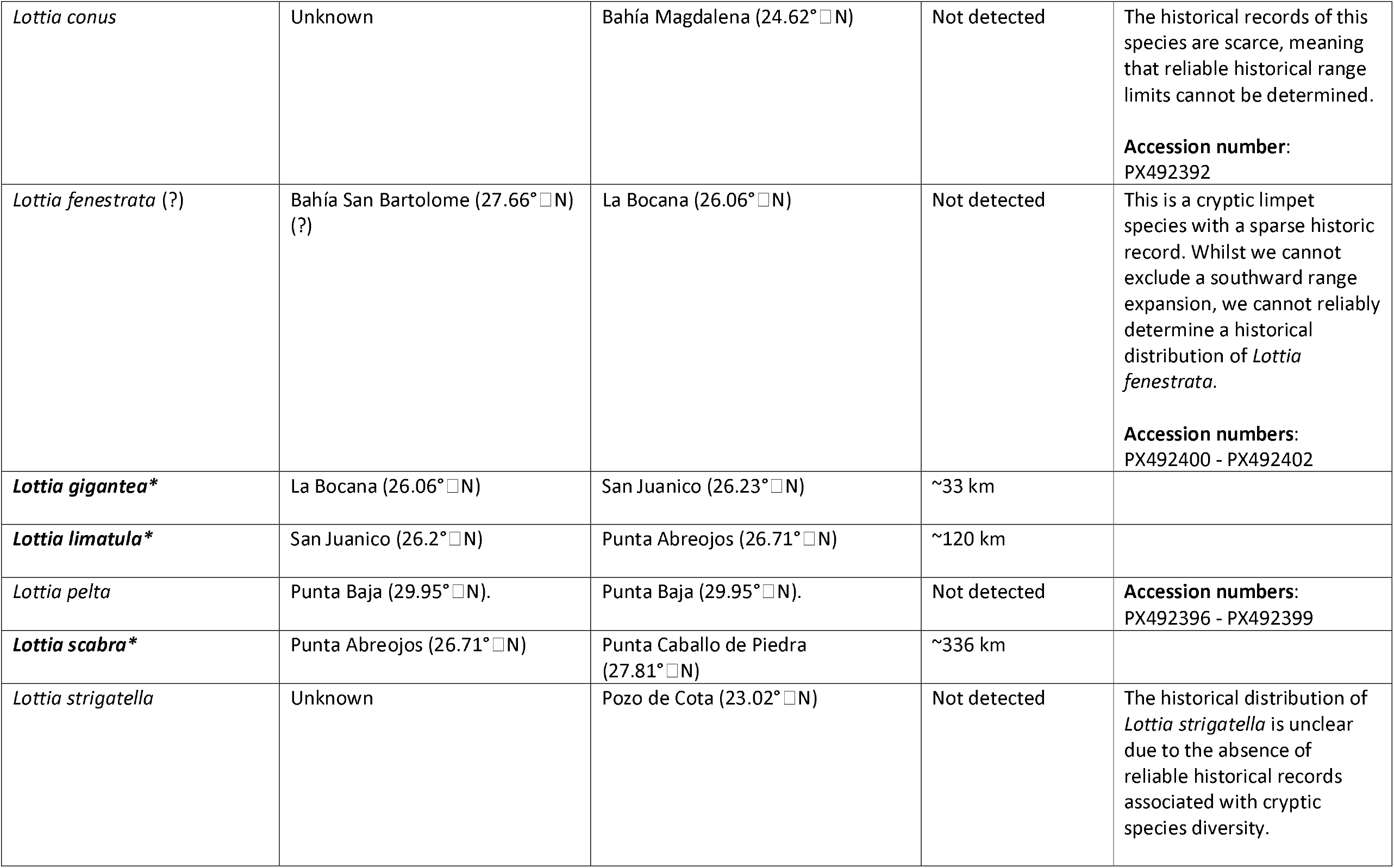
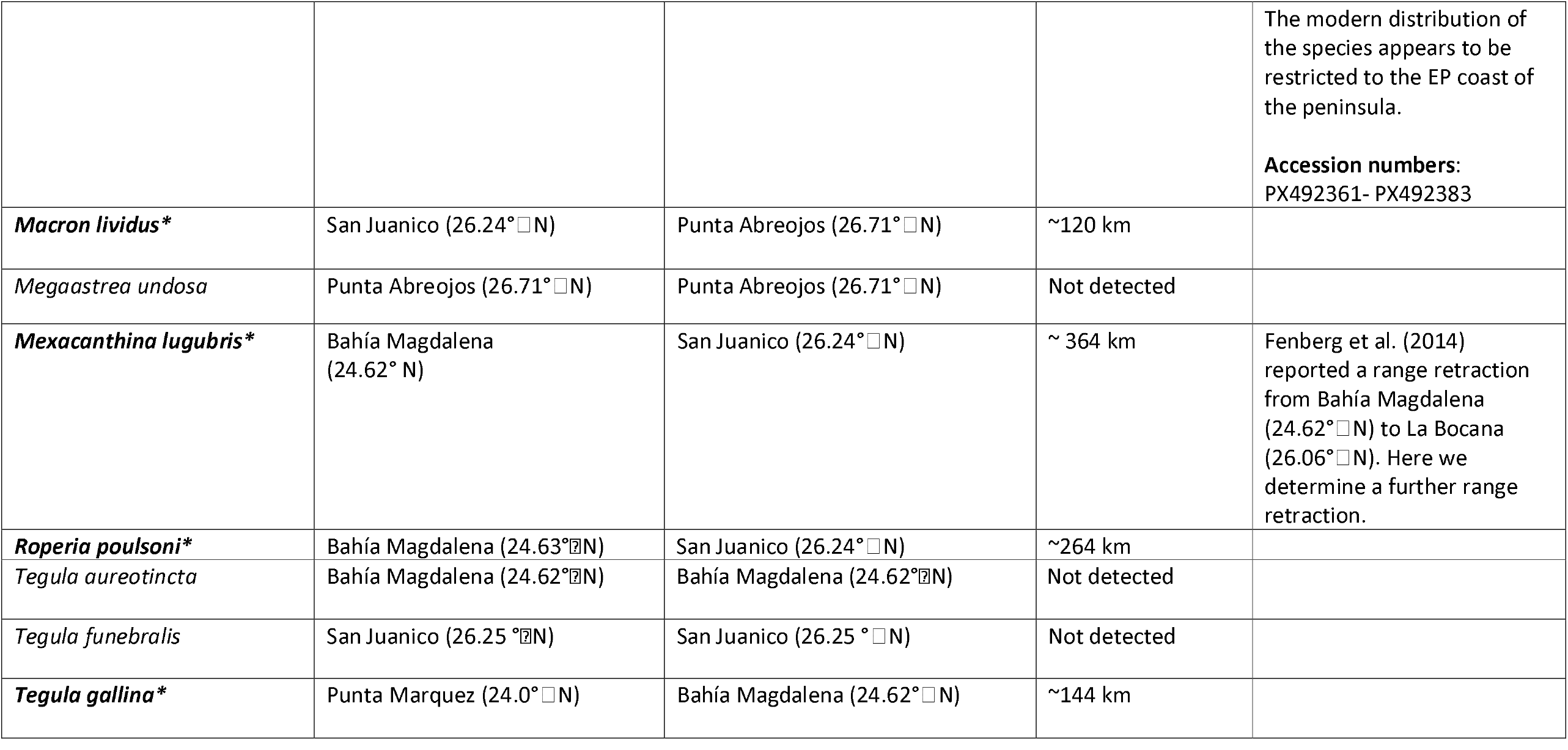
Modern and historical southern range limits and range shifts in km of temperate rocky shore invertebrates. Range retracting species are bolded and marked with *. Detailed description of species ranges, and relevant historic species records are described in Supplementary Information S1. Where applicable we provide the Accession Numbers to the partial COI genetic markers obtained in the study.

**Figure 2.**
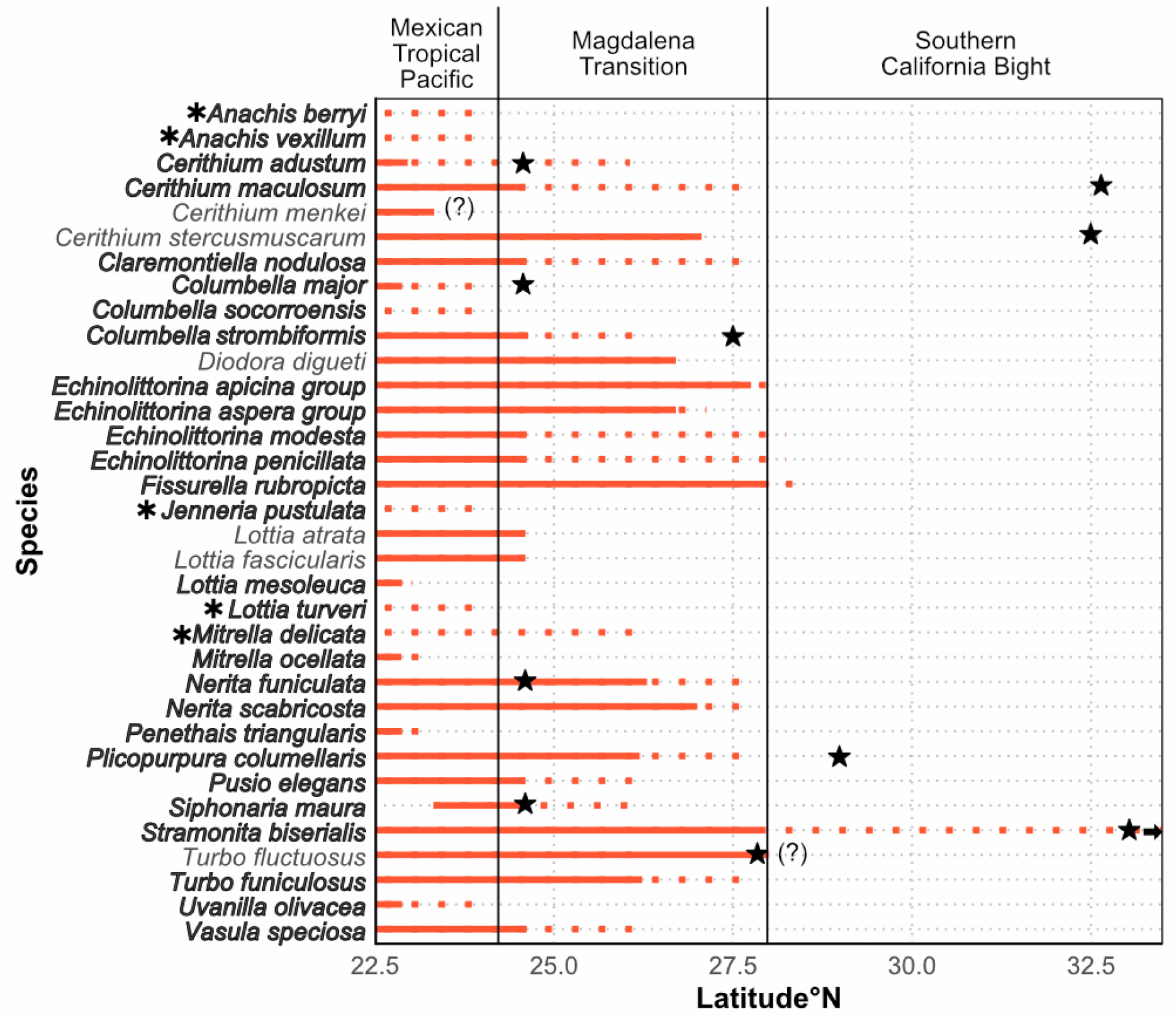
Tropical and subtropical species detected along the Eastern Pacific coastline of Baja California. Species restricted to the Gulf of California have been excluded from the figure. Solid lines represent historic range extent at the northern edge of the species range. Bolded species represent range expanding species, with the extent of the recent range expansion marked with dashed lines. For *Stramonita biserialis*, the arrow indicates the modern range extent extends beyond the latitudinal limit on the figure. Stars indicate the northernmost fossil records of the corresponding species, dated to the Pleistocene epoch (1.8 million to 10,000 years ago). (?) indicates uncertainty regarding historic or present range (Table 1).

**Figure 3.**
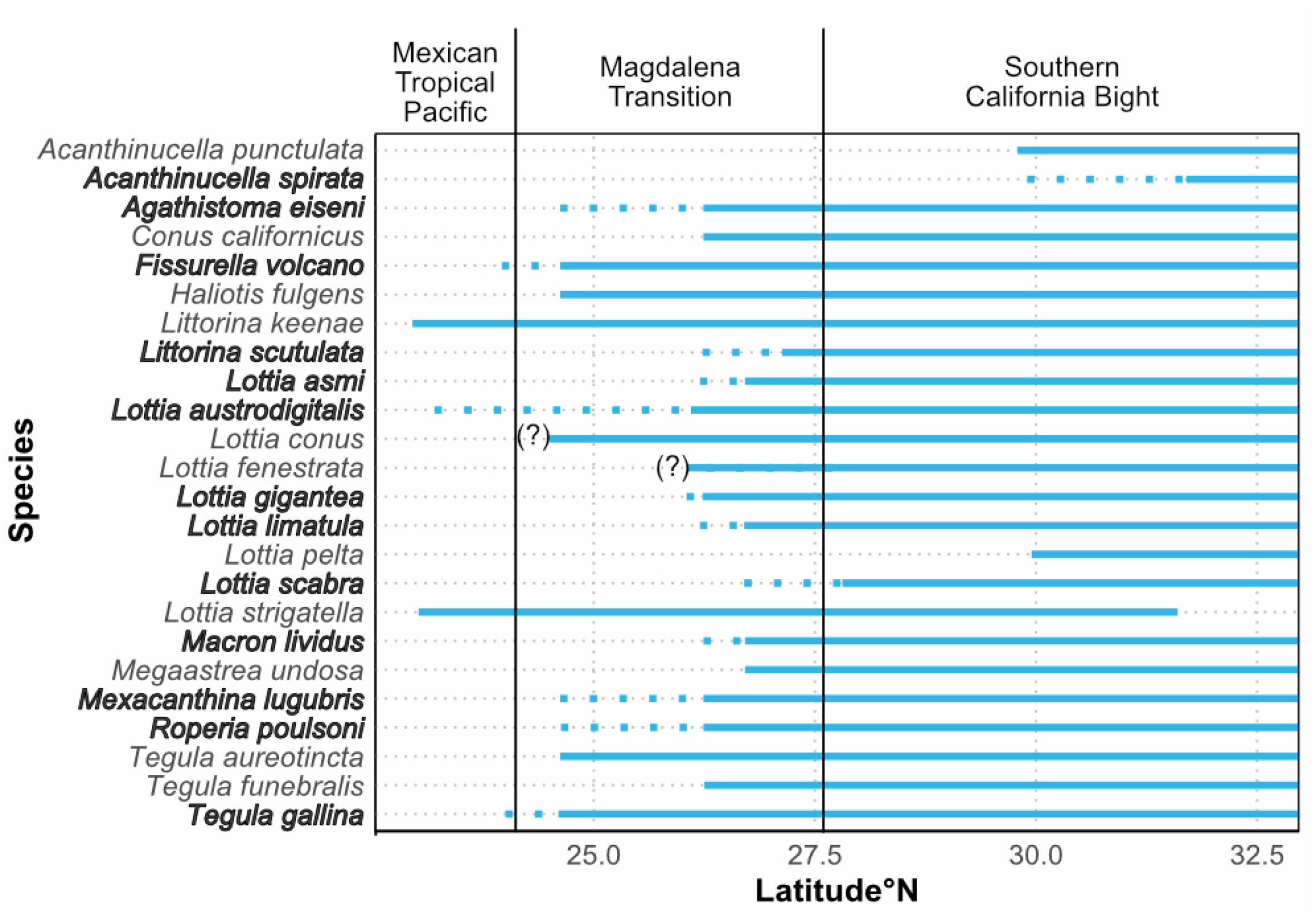
Temperate species detected along the EP coastline of Baja California. Species restricted to the GoC have been excluded from the figure. Solid lines represent the modern range extent at the southern, trailing-edge of the species range. Bolded species represent range retracting species, with the extent of the recent range-retraction marked with dashed lines (i.e. historic range limit). (?) indicates uncertainty regarding historic or present range (Table 2).

We found a number of range expanding species that have previously been restricted to the GoC but have now expanded to the EP coast. Those species include *Anachis berryi, A. vexillum, Jenneria pustulata, Lottia turveri* and *Mitrella delicata*. Meanwhile, *Columbella socorroensis* which was historically not present on the Baja California Peninsula, and might have expanded from Socorro Island, Mexico (18.85° N), although the historic record of this species is scarce.

### Pleistocene records of range expanding tropical species

We obtained Pleistocene fossil records for eight range expanding tropical species: *Cerithium adustum, C. maculosum, Columbella major, C. strombiformis, Nerita funiculata, Plicopurpura columellaris, Siphonaria maura* and *Stramonita biserialis* (Table 1; Figure 2). For *C. adustum, C. maculosum, C. major, C. strombiformis, P. columellaris* and *S. biserialis*, the northernmost recorded Pleistocene fossil has been found beyond the modern range of the species. For *N. funiculata*, the northernmost recorded Pleistocene fossil comes from Bahía Magdalena (24.6° N) which falls well within the historic range of this species. For *S. maura*, the northernmost fossil record comes from its historic northern range limit which is also at Bahía Magdalena (24.6° N), although we report a range expansion of this species beyond this point. We also identified Pleistocene records for two tropical species which are not currently experiencing a range expansion; *C. stercusmuscarum* and *Turbo fluctuosus*. Whilst the northernmost fossil record of *T. fluctuosus* comes from the same location as its historic and modern range limit, the Pleistocene fossil for *C. stercusmuscarum* comes from well beyond the current range limit of this species (Figure 2).

### *Range retraction and decreases in abundance of* Lottia gigantea

We detected a trailing-edge range retraction for *L. gigantea* of 33 km from La Bocana (26.06° N) to San Juanico (26.24° N). Due to the local extinction of this species at La Bocana, we were able to obtain size data for the limpets at Santa Rosalillita, Punta Abreojos and San Juanico only (Figure 4). Consequently, La Bocana was excluded from the statistical analysis, but the median shell size of *L. gigantea* at this site in 2003-2006 sampling period was 32.5 mm (IQR = 9, n =68). At San Juanico, median shell size increased from 28 mm (IQR = 13, n = 86) in 2003-2006, to 41 mm (IQR = 20, n = 19) = in 2022-2024 (Mann–Whitney U test, Bonferroni-adjusted p < 0.001). At Punta Abreojos, median sizes decreased from 25 mm (IQR = 17, n = 198) in 2003-2006, to 20 mm (IQR = 12, n = 55), in 2022-2024 (Mann–Whitney U test, Bonferroni-adjusted p < 0.05). At Santa Rosalillita, the median shell size decreased from 36 mm (IQR = 16, n = 107) to 28 mm (IQR = 16, n = 101) in 2022-2024 (Mann– Whitney U test, Bonferroni-adjusted p < 0.001).

**Figure 4.**
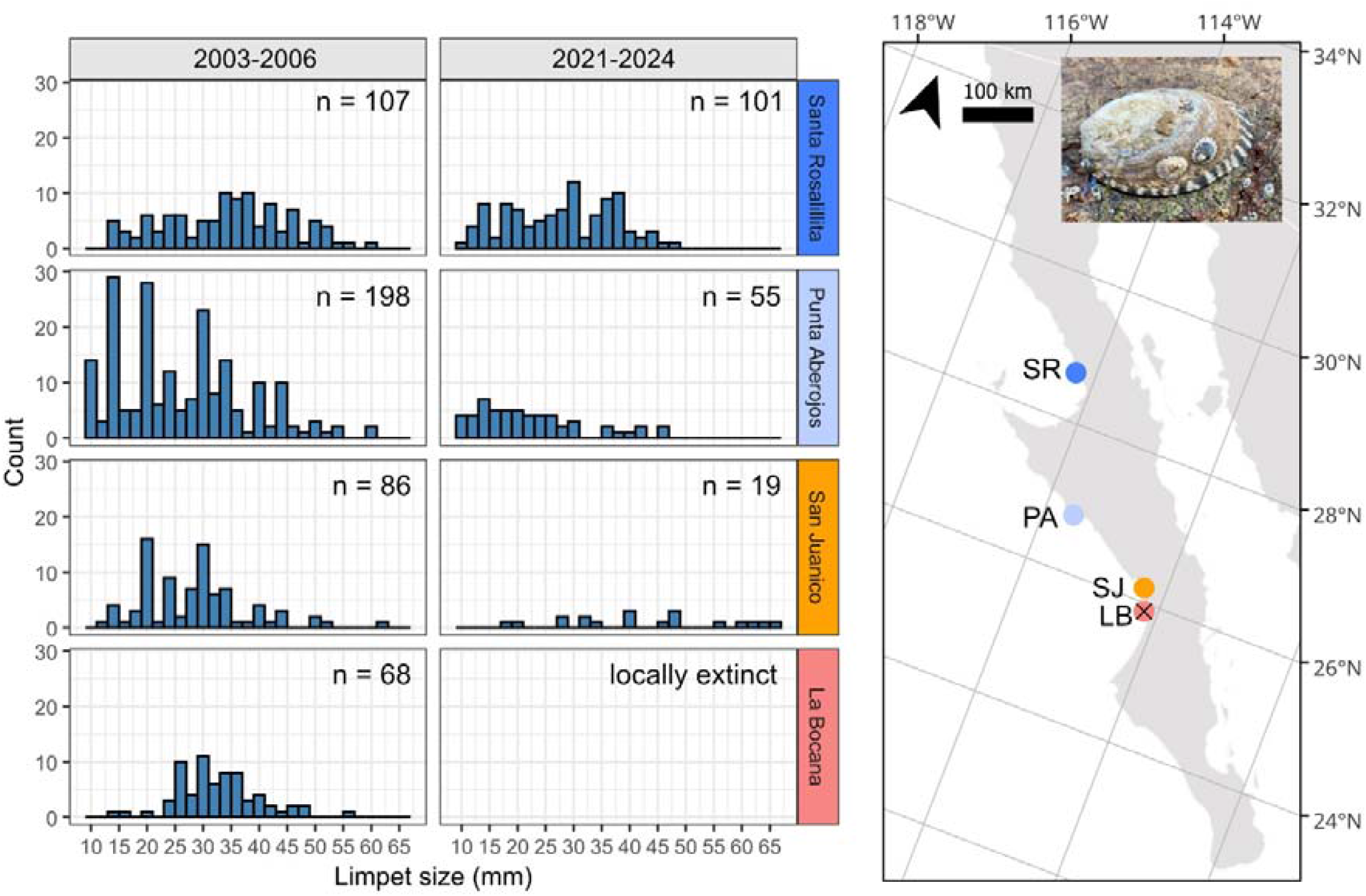
Size-frequency distribution of *Lottia gigantea* shell size during historic surveys conducted between 2003-2006, and during recent surveys conducted between 2021-2024. The limpet size and frequency are reported for the southern range-edge of the species. The size-frequency distributions indicate a local extinction at La Bocana (LB), and a potential reduction in recruitment at San Juanico (SJ) and Punta Abreojos (PA) as demonstrated by reduced number of small individuals during modern surveys. The size-frequency distribution remains largely unchanged at Santa Rosalillita (SR).

## Discussion

### Patterns of tropicalisation

Documenting tropicalisation is often challenging as the historic distribution data needed to detect range shifts are often unavailable (Klaassen et al., 2025; McClenachan et al., 2012; Przeslawski et al., 2012; Zarzyczny et al., 2024b), and modern biodiversity surveys are labour intensive (Borges et al., 2023; Ross et al., 2025), and rely on taxonomic expertise (Hayes et al., 2025; Zarzyczny et al., 2025). Here, we combined historical surveys, museum collections and online biodiversity databases with modern field surveys to document distributional changes of 32 gastropod species across the Baja California coastline. Whilst tropicalisation continues to be documented in a range of marine systems globally (Zarzyczny et al., 2024a), our research represents one of the first comprehensive tropicalisation studies across intertidal rocky shores. Moreover, we demonstrate that these shifts are occurring across established biogeographic boundaries (Blanchette et al., 2008; Fenberg et al., 2015; Spalding et al., 2007).

We detected range expansions of multiple tropical gastropods along the EP coastline, and to a lesser extend within the GoC. Although the number of study sites within the GoC was limited, the pattern of tropical incursions along the EP coastline is robust and consistent with the pattern of tropicalisation. Moreover, several species which historically have been restricted to the GoC (*A. berryi, A. vexillum, J. pustulata, L. turveri*, and *M. delicata*), appear to have expanded their ranges to the EP coastline, progressing across established biogeographic boundaries. Similarly, some tropical/subtropical gastropods are range extending beyond biogeographic boundaries on the EP coastline. *Cerithium adustum* which was previously restricted to the Mexican Tropical Pacific, is now present within the Magdalena Transition biogeographic region. Meanwhile, *S. biserialis* and *F. rubropicta* have extended their ranges into the temperate Southern California Bight region.

Distinguishing between vagrant tropical species, and range extending taxa is critical when interpreting climate-induced range shifts (Bates et al., 2014; Zarzyczny et al., 2024a). Vagrant species are typically represented by sporadic records (Bates et al., 2014), and their persistence is restricted by winter temperatures that inhibit overwinter survival and reproduction (Booth et al., 2007; Figueira & Booth, 2010). As our surveys were conducted primarily during winter months, and multiple sites were revisited repeatedly between 2017 and 2024, we are confident that the species documented here represent genuine range expansions, rather than transient occurrences.

In addition to range expansion, we also identified several cases of range retractions among warm-temperate species. Such contractions are typically more difficult to confirm than expansions, as multiple surveys or complementary lines of evidence are required to verify local extinctions (Poloczanska et al., 2013; Zarzyczny et al., 2025; Zarzyczny et al., 2024a). Early stages of range retractions are often detectable through reduced ecological performance or declining abundance prior to local extinction (Bates et al., 2014), and are likely associated with increased stress at the trailing-edge of the species’ range (King et al., 2020). This pattern is clearly demonstrated by *L. gigantea;* although we confirmed its southern range limit has shifted northwards by ∼33 km, our size-frequency analysis indicates significant demographic effects well-beyond the current range boundary. At San Juanico and Punta Abreojos, abundances have declined by approximately 72-77% compared to historical baselines. Additionally, both sites demonstrate marked losses of smaller individuals indicating a substantial reduction of recruitment. The near absence of juveniles at San Juanico suggests that recruitment failure is already underway. If these trends persist, further range retraction of *L. gigantea* appears likely in the near future.

Range retractions are also occurring across biogeographic boundaries. For example, the trailing-edge limit of *F. volcano, L. austrodigitalis*, and *T. gallina* historically extended into the Mexican Tropical Pacific. However, all three species have now retracted their ranges into Magdalena Transition. Additionally, *L. scabra* was previously reported to extended south into Magdalena Transition and has since retracted its range into the Southern California Bight. However, given the morphological similarities between *L. scabra* and a more southern species, *L. conus* this observation should be interpreted with caution (Supplementary Information S1).

The observed range shifts may indicate that established biogeographic boundaries along the Baja California coastline are beginning to erode. Several tropical and subtropical taxa are now extending beyond biogeographic limits, suggesting that long-standing environmental and oceanographic filters (Fenberg et al., 2015) are becoming increasingly more permeable. In contrast, temperate species are gradually becoming lost from entire biogeographic regions, highlighting an asymmetry in how range expanding and range retracting species are responding to warming (Poloczanska et al., 2013). These opposing dynamics are reshaping coastal community structure, potentially weakening biogeographic partitioning, and will likely have important ecological and evolutionary consequences.

### Ecological and evolutionary consequences of tropicalisation

The redistribution of species along the Baja California coastline is generating novel community compositions which are likely leading to new species interactions. As tropical and subtropical taxa establish in new regions, predator-prey dynamics (Fenberg et al., 2023), competition (Steger et al., 2021) and other interspecific relationships can change (Minguito-Frutos et al., 2025; Vergés et al., 2016). Such changes have the potential to alter structure and functioning, but also evolutionary trajectories of species living on rocky shore ecosystems.

While functional ecology assessments are beyond the scope of this study, some the findings included can provide insights into species traits that may influence tropicalisation dynamics. Many range extending species possess traits that facilitate dispersal and colonisation (Coni et al., 2022; Downie et al., 2025). These taxa appear to be more generalist in their habitat and dietary preferences and exhibit high dispersal capabilities which likely underpin their ability to extend ranges across biogeographic boundaries (Downie et al., 2025; Zarzyczny et al., 2024a). For example, *S. biserialis* is proving to be a particularly successful range-extending species (Figure 2; Table 1). *Stramonita biserialis* is a generalist predator, capable of feeding on a range of intertidal species including bivalves, other gastropods, and barnacles (Fenberg et al., 2023; Herbert, 2004). This species also has a high dispersal potential given Its long pelagic larval duration of over 2 months (Claremont et al., 2011; Fenberg et al., 2023) likely influencing its successful range expansion.

Moreover, as species interactions are thought to be stronger in the tropics in comparison to temperate regions (Freestone et al., 2021; Schemske et al., 2009), arrival of tropical and subtropical species may therefore intensify ecological interactions in newly colonised communities. Predation pressure in particular, is widely recognised to decline with increasing latitude (Ashton et al., 2022; Freestone et al., 2021). Tropical ecosystems support higher predation rates, greater diversity of predators, and predators of larger body size (Ashton et al., 2022; Freestone et al., 2021). Consequently, range-expanding tropical predators may exert substantial impacts on temperate prey. For example, predation by range-expanding tropical predators is thought to induce morphological defences in the temperate barnacle, *Tetraclita rubescens*, producing bent shell thought to reduce predation risk (Fenberg et al., 2023). Similarly, the tropical crown-of-thorns star fish, *Acanthaster cf. solaris*, which typically does not predate on coral within its native range, has been observed preying on endemic and high latitude corals within its recently expanded range along the western Pacific coast of Australia (Sommer et al., 2025). Similarly, outbreaks of- and mass predation by *A. cf. solaris* (Rodríguez-Villalobos & Ayala-Bocos, 2021) and corallivore gastropod *Jenneria pustulata* (Paz-García et al., 2012) have been observed in the southern and northern Gulf of California, respectively. Such observations are particularly problematic given the slow-growing nature, and limited recruitment of the high-latitude corals (Sommer et al., 2024, 2025). Here, we have documented the range expansions of several charismatic tropical gastropods including *Claremontiella nodulosa, Penethais triangularis, Plicopurpura columellaris, Vasula speciosa*, and the previously mentioned *S. biserialis* and *J. pustulata*. Whilst predation dynamics under tropicalisation remain relatively understudied (Zarzyczny et al., 2024a), the few examples of above highlight how these range-expanding tropical predators can impact temperate ecosystems by altering trophic interactions.

In contrast to range-extending taxa, temperate species undergoing range retraction may be at risk of losing genetically distinct trailing-edge populations (Nielsen et al., 2024; Zarzyczny et al., 2024b; Zarzyczny et al., 2024a). Genetic structuring in many temperate marine taxa indicates that southern populations frequently harbour unique haplotypes, separated from northern populations by long-standing phylogeographic breaks (Nielsen et al., 2024; Riginos & Nachman, 2001; Zarzyczny et al., 2024b). For instance, *L. gigantea* which has undergone a range retraction in recent years (Figure 3), has private haplotypes (i.e. unique genetic diversity) within its trailing-edge populations (Nielsen et al., 2024). Other temperate limpets in this region, such as *L. strigatella* and *L. conus*, also possess genetically distinct southern populations (Zarzyczny et al., 2024b). Given that many of the temperate species (in particular limpets) share comparable dispersal capabilities and are subject to similar barriers to gene flow (Zarzyczny et al., 2024b), it is possible that other range retracting species identified in our study also harbour unique genetic diversity within their southern populations. The continued retraction of these populations could lead to irreversible loss of unique evolutionary lineages. Such losses are particularly concerning given that reduced intraspecific genetic diversity could lead to decreased adaptive potential of species (Hudson et al., 2021; Willi et al., 2006).

### What can the Pleistocene fossil record tell us about tropicalisation patterns?

Range expansions have occurred repeatedly in the past (Hellberg et al., 2001; Marko et al., 2010). Fossil evidence demonstrates that several tropical gastropod species previously extended well beyond their historic and even modern distributions in the Pleistocene. However, given the rate of ocean warming under the contemporary climate change is much faster than during Pleistocene (Intergovernmental Panel On Climate Change, 2023), the pace of modern range shifts is unprecedented. Moreover, whilst during the Pleistocene climatic fluctuations species retreated towards the equator during cooler periods (Bagley et al., 2013; Hofreiter & Stewart, 2009), future climatic trajectories are unknown, and it is impossible to tell whether contemporary expansions will be followed by similar reversals.

Furthermore, as some of the range-extending species have occupied latitudes beyond their present ranges during the Pleistocene warm periods (Figure 2), they may share evolutionary history with resident temperate taxa (Fenberg et al., 2023; Zarzyczny et al., 2024a). Such historical overlap could have two consequences. On one hand, range-extending species may retain adaptations or traits that enable them to successfully colonise higher latitudes as climatic envelope expands (Hill et al., 2011), giving them an advantage over resident competitors or prey. On the other hand, some degree or adaptation or ecological familiarity could modulate interactions between the range expanding and resident species, potentially reducing the novelty of these ecological encounters (Mech et al., 2019; Zarzyczny et al., 2024a).

### Challenges and future directions

Tropicalisation is occurring globally across diverse ecosystems (Zarzyczny et al., 2024a), yet intertidal rocky shores have been largely overlooked. Our study provides new evidence that these habitats are also experiencing climate-driven shifts in species composition. Despite growing tropicalisation research, many regions and ecosystems remain understudied. Continued documentation is essential to guide conservation and management decisions, particularly given the ecological and evolutionary consequences of novel species arrivals and temperate species losses (Vergés et al., 2016; Zarzyczny et al., 2024a).

Accurate documentation of tropicalisation relies on species records spanning large spatial and temporal scales (Zarzyczny et al., 2024a). Historical surveys and museum collections used to establish historic species ranges are often sparse, subject to regional and taxonomic bias, and incomplete records (Meineke & Daru, 2021; Parker et al., 2024; Zarzyczny et al., 2024b). Taxonomic misidentifications, cryptic species and ongoing taxonomic reassignments further complicate range interpretations. Emerging approaches, such as the use of computer vision (application of artificial intelligence to pattern recognition) can help detect inaccuracies in museum records (Hollister et al., 2025) and offer a promising means to address some of these limitations.

Cryptic diversity remains a major challenge for documenting tropicalisation in contemporary assemblages, particularly among invertebrates (Shin & Allmon, 2023). Many gastropod species share superficially similar morphotypes and large intraspecific variation, making it difficult to resolve species boundaries without molecular data (Hollister et al., 2023; Williams et al., 2012). Currently, the most effective means of resolving such lineages involves genetic barcoding, which typically requires collecting, and sacrificing individuals for DNA extraction (Shin & Allmon, 2023). While this approach can provide robust taxonomic resolution, it is labour-intensive, and often impractical for large-scale monitoring. The use of environmental DNA (eDNA) data, which does not rely on collection of individual specimens and sequencing but rather examines community sequence data from DNA shed by organisms into the local environment, is being explored as an alternate approach for monitoring range shifting species (Holman et al., 2022; Zarzyczny et al., 2025). Furthermore, emerging techniques using artificial intelligence offer promising, non-destructive alternatives. For example, machine learning models can be trained on shell photographs to accurately classify species under controlled conditions (Hollister et al., 2023) and predict environmental conditions for species distributions (Bolaños-Durán et al., 2025). With continued advances, these models may eventually enable identification of individuals in situ from field images, reducing reliance on destructive sampling, improve species identification and predict environmental conditions for shift in species distributions under the effects of tropicalisation.

## Conclusions

Our findings demonstrate that tropicalisation is actively reshaping rocky shore assemblages along the Baja California peninsula, with tropical and subtropical species expanding across long-standing biogeographic boundaries, and temperate taxa undergoing pronounced retractions. Continued documentation of these dynamics is essential for detecting early ecological and evolutionary consequences of tropicalisation. Moving forward, robust monitoring efforts which integrate historical distribution data, modern field surveys, along with molecular and computational tools will be crucial for improving our understanding of these processes, particularly in understudied regions and taxa which harbour cryptic diversity.

## Acknowledgements

We thank Lindsey T. Groves (Natural History Museum of Los Angeles), Dr. Henry W. Chaney and Dr Daniel L. Geiger (Santa Barbara Museum of Natural History), John Slapcinsky (University of Florida Museum), Paul Callomon and Dr Gary Rosenberg (Academy of Natural Sciences of Drexel University, Philadelphia), Prof. Jerry Harasewych (Smithsonian National Museum of Natural History), Johanna Loacker (California Academy of Sciences), and Alex Kittle (Delaware Museum of Nature and Science) for providing their support in accessing collection images and sharing their knowledge which allowed us to confirm species occurrences.

We thank Erick X. Treviño Balandra and Eva Gallardo for their support during fieldwork, and Fabian Aspey-Gay for assisting with some of the DNA extractions. Finally, this work was supported by an Early Career Research Grant from the Malacological Society of London, and a Heredity Fieldwork Grant from the Genetics Society to KMZ; the Natural Environmental Research Council (NE/S007210/1) to KMZ and (NE/X011518/1) to PBF; and The Royal Society (RG2017R1) to PBF. The permit for sample collection was provided by the Secretaría de Agricultura, Ganadería, Desarrollo Rural, Pesca y Alimentación (SAGARPA, Permiso de Pesca de Fomento No. PPF/DGOPA-291/17 and PPF/DGOPA-010/19).

## Data Availability Statement

Species detection data is available in the Supplementary Information. Partial COI genetic markers used to barcode cryptic species are available on Genbank (https://www.ncbi.nlm.nih.gov/genbank/) with accession numbers PX492361-PX492409.

