## Supplementary Information for "Tropicalisation of rocky shore gastropod communities along the Baja California Peninsula"

KMZ: 0000-0003-1675-2090

DAPG: 0000-0002-1228-5221

MR: 0000-0002-2195-6605

STW: 0000-0003-2995-5823

PBF: 0000-0003-4474-176X

**Supplementary Information S1: Species Accounts**

This document contains summaries of changes in geographic distribution of the studied species between 2017-2024, in comparison to their historical ranges obtained from the literature and museum collections (typically year 2000 and earlier, unless stated otherwise).

**Abbreviations:**

Collections and museums

**ANSP:** Academy of Natural Sciences

**CAS**: California Academy of Sciences

**LACM**: Los Angeles Country Natural History Museum

**PRI**: Paleontological Research Institution

**SBNMH:** Santa Barbara Museum of Natural History

**USNM**: Smithsonian Institution, National Museum of Natural History.

General

**EP:** Eastern Pacific

**GoC:** Gulf of California

**Species detected during the study (alphabetical).**

***Acanthinucella punctulata***

The southernmost historic record of *Acanthinucella punctulata* appears to be from Isla San Geronimo (29.79° N; SBMNH catalogue number 113551; Chaney et al., 2024a), and Rancho Socorro (30.33° N; LACM 1966-3.20; Groves and Mertz, 2025a) on the EP coast. Whilst our surveys were limited to the main coastline, we detected *A. punctulata* at the nearest site to the historic range limit, at La Chorera (30.47° N) suggesting the southern range of this species likely remains unchanged.

***Acanthinucella spirata***

The historic southern range limit of *Acanthinucella spirata* is reported to be Punta Baja (22.9° N; Flagor and Bourdeau, 2018). We did not detect this species at Punta Baja during our surveys in 2021-2022, or 2024. We place the current southern range limit of *A. spirata* as Campo Kennedy, at Punta Banda (31.70° N), and report a range retraction of ~288 km. Interestingly, whilst we report a range retraction of the southern range limit of this species, Flagor and Bourdeau (2018) have reported a recent range expansion at the northern range limit in California. The combined observations suggest a clear poleward range shift experienced by this this species.

***Agathistoma cortezianum***

Historically distributed throughout the GoC, from the head of the Gulf, to Cerralvo Island, southeastern GoC and to Guayamas, Sonora. We detected *Agathistoma cortezianum* at La Gringa (29.05 ° N), and Agua Verde (25.52 ° N) but not La Paz, which corresponds to the historic records of this species.

***Agathistoma eiseni***

Historically, *Agathistoma eiseni* was distributed from Bahía Magdalena (24.62 ° N) to Los Angeles, California (34° N; Hellberg, 1998; Keen 1971). We have not detected *A. eiseni* south of San Juanico (26.24° N) suggesting a range retraction of 264 km.

***Anachis berryi***

Historically restricted to El Pulmo Reef (23.43° N) located at the tip of the Baja California Peninsula, found at 1 to 3m depth (Keen, 1971). Whilst we surveyed the intertidal region only, we detected *Anachis berryi* at Punta Marquez (23.96° N), suggesting a possible range expansion of at least 264 km.

***Anachis vexillum***

Historically restricted to the GoC, from the head of the Gulf, south to Acapulco (Keen, 1971). We detected individuals of *Anachi vexillum* within the GoC at La Gringa (29.05° N) and on the outer EP coastline at Punta Marquez (23.96° N), suggesting a potential range expansion of ~264 km.

***Callianax biplicata***

Historically, the distribution of *Callianax biplicata* ranges from Vancouver Island (48-51° N; Abbott, 1954), south to Bahía Magdalena (24.6° N; SBMNH 123253; Chaney et al., 2024). We detected this species at Punta Abreojos (26.71° N) only. However, as *C. biplicata* typically inhabits sandy substrates in the intertidal or shallow tidal zones of the EP coast (Kelly et al., 2021), our surveys did not capture its true modern distribution.

***Cerithium adustum***

According to Keen (1971), the distribution of this species ranged from the GoC, South to Peru. Two dry specimens (catalogue number 154666) preserved at the Santa Barbara Museum of Natural History, have been collected at El Tulo (22.96° N), at the southern tip of Baja California Sur, in 1964 (Chaney et al., 2024). We have detected two populations further north on the Eastern Pacific Coast, one at Punta Marquez (23.96° N) and one as far as La Bocana (26.06 ° N), suggesting a range expansion of ~552 km, for this species.

A Pleistocene fossil specimen of this species, collected in 1925 at Bahía Magdalena (24.6 ° N; CASG:FOSSIL:84663), is catalogued at California Academy of Sciences (Roopnarine and Garcia, 2024). This northern record could be associated with the climatic fluctuations during the Pleistocene period (Hofreiter and Stewart, 2009), leading to population range expansions or shifts into northern refugia, subsequent range retraction during cooler periods, and a recent range expansion over the past 60 years.

***Cerithium maculosum***

The historical distribution of this species ranged from Bahía Magdalena (24.6 ° N), through the GoC and south to Mazatlan (Keen, 1971). Since, this species has expanded its range along the eastern Pacific Coastline, with populations present at San Juanico (26.23 ° N), Punta Clambey (27.65 ° N), and as far north as Punta Caballo de Piedra (27.81 ° N), suggesting a range expansion of ~720 km.

***Cerithium menkei***

Whilst Keen (1971) places the northern distribution of *Cerithium menkei* as being limited to the GoC, specimens from the EP coast have since been identified and databased at the LACM. The six lots of specimens were collected after the Keen (1971) publication. The three lots (LACM 1979-26.19, LACM 1979-25.21 and LACM 1979-24.11) have been collected from Punta Marquez (23.95° N), Punta Pequena (26.23° N), and Punta Santo Domingo (26.32° N) in 1979, respectively. Additionally, three lots of specimens have been collected from Punta Pequena in 1971 (LACM 1971-6.15, LACM 1971-179.6 and LACM 1971-181.16).

Interestingly, we have verified two more lots of *C. menkei* collected in 1951 all the way from the vicinity of Punta Eugenia (LACM 1951-42.8 from the east side of Punta Eugenia, 27.83 ° N; and LACM 1951-45.16 from Caleta Hancock, ~ 20 miles from Punta Eugenia, 27.81° N). These lots were not catalogued in the LACM Malacology Collection at the time of the Keen (1971) publication, therefore we do not know whether this material was examined at the time. However, all the collections were identified and verified by the expert curator, Jim McLean, in 1991 when they were databased, and during this study, we have confirmed their identification with the support from Lindsey Groves (Senior Collections Manager, LACM). Consequently, this places the northern historical range limit of *C. menkei* as far as Punta Eugenia (27.83 ° N). However, our surveys have not detected *C. menkei* on the EP Coast, north of Cerritos Point (23.33° N). It is possible that this species has a discontinuous distribution, or that it is found subtidally on the EP coastline.

***Cerithium stercusmuscarum***

Keen (1971) describes the distribution of *Cerithium stercusmuscarum* as “Baja California to Peru”. Northernmost historic records of this species are from San Ignacio Lagoon (27.06° N; ANSP Catalogue number 17756; Rosenberg, 2025). We detected *C. stercusmuscarum* along the EP coast of Baja California, with the northernmost record of this species at San Juanico (26.24° N). We did not detect this species at Punta Abreojos (26.71° N) which although latitudinally is located south of San Ignacio Lagoon, this site is further west along the EP coastline, and hence, falls outside of this species historic range. Although we did not survey San Ignacio Lagoon (27.06° N), we have confirmed modern records of this species in this area using iNaturalist (Axley, 2025; Paredes Montesinos, 2024).

Interestingly, fossil specimens dating back to Pliocene and Pleistocene (PRI Catalogue Number 45331) have been collected from the northern portion of Baja California near Tijuana (32.4° N) (Skibinski, 2025). Several Quaternary fossil specimens have also been identified from Punta Eugenia area (27.8° N; Randall, 2021). Whilst we have not detected a recent northward range expansion of this species, such northern records could be associated with the climatic fluctuations during the Pleistocene period (Hofreiter and Stewart, 2009), leading to population range expansions or shifts into northern refugia, and subsequent range retraction during cooler periods leading to the present geographic distribution.

***Claremontiella nodulosa***

The historic distribution of *Claremontiella nodulosa* ranges from Bahía Magdalena (24.62° N), through the GoC and south to Guyamas (Keen, 1971). We detected this species at all of our surveyed sites within the GoC, and at multiple sites along the EP coast. On the EP coast, we place the modern northern range limit of *C. nodulosa* as Punta Caballo de Piedra (27.81° N), demonstrating a recent range expansion on 720 km.

***Columbella aureomexicana***

According to Keen (1971), *Columbella aureomexicana* is historically distributed from Cedros Island (28.2° N), Baja California through the GoC and down to Sonora. We have identified preserved specimen (LACM 1983-146.1) from El Tomatal (28.48 ° N) collected in 1983 is stored in the LACM collections placing the historic northern range limit slightly further north than Keen (1971). Despite this, during our surveys we only detected individuals within the GoC. This finding can be attributed to the fact that many *Collumbella* spp. including *C. aureomexicana* are found in subtidal waters (for example SDNHM catalogue number 51746; San Diego Natural History Museum 2025).

Interestingly, fossils of *C. aureomexicana* dating back to the Pleistocene, have been identified from as far north as Newport Beach, California (33.6° N; LACMIP 66.1050 and LACMIP 22539.71; Hendy and Hook, 2024), suggesting a more northern range limit, likely associated with the climatic fluctuations during the Pleistocene period (Hofreiter and Stewart, 2009), leading to population range expansions or shifts into northern refugia, and subsequent range retraction during cooler periods leading to the present geographic distribution.

***Columbella fuscata***

The historical distribution of this snail extends from Peru, through the GoC and north to Bahía Magdalena Bay (24.6° N) on the EP coast (Keen, 1971). A single preserved specimen from Bahía Magdalena is catalogued at the ANSP (cat no. 152162) albeit with no collection date. Another specimen from Bahía Magdalena is a Pleistocene fossil collected in 1925, which is held at CAS (CASG:FOSSIL:85914). Four preserved specimens from near Punta Conejo (24.25 ° N), albeit with no date are held in the SBMNH collections (cat. no. 366353). Northernmost dated (1976 – 1984) specimens from the EP coast are from Cabo San Lucas (22.90 ° N). However, we have not detected this species north of Punta Marquez (23.96 ° N), which can be attributed to the fact that many *Collumbella* spp. including *C. fuscata* are found in subtidal waters (e.g. SDNHM Catalogue number 52015; San Diego Natural History Museum, 2025), which fall beyond the scope of our intertidal surveys.

***Columbella major***

Southern portion of the GoC, down to Peru (Keen, 1971), with a single specimen from Cabo San Lucas (22.88 ° N) collected in 1984 on the EP coast (cat no. 237474; Delaware Museum of Nature and Science, 2025). We have observed this species as far north as Punta Marquez (23.96 ° N).

***Columbella socorroensis***

Records of historical specimens are very rare, with scarce records from the Socorro Island, Mexico (Keen, 1971). However, we have observed this snail at Punta Marquez (23.96 ° N), suggesting a potential range expansion to the Baja California Peninsula of ~564 km.

***Columbella strombiformis***

Historically present throughout the Gulf of California and south to Peru (Keen, 1971). A single undated record from Bahía Magdalena (24.63° N) is held in the ANSP collection (Catalogue number 152163; Rosenberg, 2025). Albeit undated, this preserved specimen was collected by Orcutt, C.R. (1864 – 1929), and was likely collected in 1917 during his visit to Bahía Magdalena (Orcutt and San Diego Society of Natural History, 1918). We detected it as far as San Juanico (26.24° N), with more sightings at Punta Marquez (23.96° N) suggesting a range expansion of 264 km from Bahía Magdalena.

A Pleistocene fossil specimen from the Punta Eugenia region is catalogued at CAS (cat no. CASG:FOSSIL:92751; Roopnarine and Garcia , 2024), suggesting a more northern distribution in the past, likely associated with the Pleistocene climatic fluctuations (Hofreiter and Stewart, 2009).

***Conus californicus***

Based on Keen’s (1971) description, *Conus californicus* appears to have a warm temperate or a cosmopolitan distribution, with its core range within southern California, and sporadic occurrences in the Panamic province. A southernmost preserved specimen (USNM 102251) was collected in 1911 and is from Ballenas Bay (26° N; Orrell, 2025). We detected this species at San Juanico (26.23 °N) suggesting the range of this species likely remains unchanged.

***Conus nux***

Historically, the northern limit of *Conus nux* is on the EP was Bahía Magdalena (24.62 °N), with its distribution extending throughout the whole GoC, and down to Ecuador (Keen, 1971). We only detected this species on the rocky shore of Punta Marquez (23.96 ° N), however, this can be explained by the fact that this species also occupies the shallow subtidal habitat (for example García-Rojas *et al.,* 2024) which we did not survey. Consequently, our surveys are unable to determine the full extent of the distribution of this species in Baja California.

***Coralliophila nux***

According to Keen (1971) the distribution of *Coralliophila nux* ranges from Baja California, south to Ecuador and Galapagos Islands. We have identified preserved specimens from the EP coast which were collected subtidally (~10m depth) from Bahía Magdalena (24.63° N) in 1977 and are catalogued at the SBMNH (cat no. 657490; Chaney et al., 2024). We have only observed this species intertidally at Cerritos Point (23.33° N) but as our surveys were restricted to the intertidal region, we cannot determine the true modern distribution of this species.

***Diodora digueti***

According to Keen (1971), *Diodora digueti* is historically known from the San Ignacio Lagoon (26.7° N) on the EP coast, with further distribution in the GoC, extending south to Ecuador. However, historic records from San Juanico (26.24° N) and Bahía Magdalena (24.5° N) are preserved at the LACM in the Malacology collection (LACM 1971-5.7, LACM 1971-181.9, LACM 1979-25.7, LACM 1971-14.22, LACM 1971-183.18) suggesting discontinuous historic distribution. We observed this species solely at San Juanico but not at Bahía Magdalena or in the GoC. Whilst we did not survey San Ignacio Lagoon, we did not observe *D. digueti* at the next two northern sites, Punta Abreojos (26.71° N) and Bahía Asunción (27.65 ° N). Consequently, we did not detect a range expansion of this species, but we are unable to determine whether a population of *D. digueti* remains at San Ignacio Lagoon.

***Echinolittorina apicina group***

We were unable to genetically separate *Echinolittorina apicina* and *E. paytensis,* using a partial COI marker. Whilst species delimitation falls beyond the scope of this study, only the distribution of *E. apicina* is expected to extend into Baja California Peninsula, whilst the northern range limit of *E. paytensis* is in Central America (Reid, 2002). According to Reid (2002), the northern range limit of *E. apicina* is within Laguna Ojo de Libre (27.75° N) in the region of Guerrero Negro. During our surveys, we detected the northern range limit of the *E. apicina* group to be Punta Morro Santo Domingo (28.24° N), suggestive of a range expansion of at least 192 km.

***Echinolittorina aspera group***

We were unable to genetically separate *Echinolittorina aspera, E. dubiosa,* and *E. tenuistriata* using a partial COI marker. Whilst species delimitation falls beyond the scope of this study, only the distribution of *E. aspera* is expected to extend to Baja California Peninsula, with the northern range limit of *E. dubiosa* and *E. tenuistriata* in Central America (Reid, 2002). According to Reid (2002), the northern range limit of *E. aspera* is located at San Ignacio Lagoon (26.7° N). During our surveys, we detected the northern range limit of the *E. aspera* group as far north as Bahía Asunción (27.13° N), suggesting a range expansion of at least 144 km

***Echinolittorina albicarinata***

Reid (2002) places the distribution of this species as far north on the EP coast as Punta Morro Santo Domingo (Laguna Manuel; 28.2° N), with further records of this species at Punta Abreojos (26.71 ° N), and Bahía Magdalena (24.62 ° N). However, during our surveys, we did not detect any *Echinolittorina albicarinata* along the EP coastline, including at Punta Morro Santo Domingo, Punta Abreojos, and Bahía Magdalena where the original specimens are collected. However, Keen (1971), describes this species as being present only on the tip of the Baja Peninsula, near Cabo San Lucas. Both authors, report the presence of *E. albicarinata* in the GoC. Whilst we did not survey the southern tip of the peninsula around Cabo San Lucas, we did observe this species at all three sampled sites within the GoC. Although we cannot exclude changes in the distribution of this species over time, our findings align more closely with the described distribution of this species by Keen (1971). However, given the discrepancies in the interpretations of this species’ historical range, we cannot draw reliable conclusions regarding potential range changes and instead provide a baseline modern range for future monitoring.

***Echinolittorina modesta***

Historically, the distribution of this species ranged from Bahía Magdalena (24.62° N) on the EP coast, to Playa Coromuel, La Paz (24.20° N) on the GoC coast of the Peninsula, and south into Central America (Reid, 2002). Within the GoC, Reid (2002) identified additional records of this species further north on Isla Carmen (~26° N), and on the mainland coast of Mexico within the Guaymas region (~27° N) (Reid, 2002). We detected *Echinolittorina modesta* as far north as Punta Morro Santo Domingo (28.24° N), suggesting a substantial range expansion of ~ 816 km on the EP coast. Within the GoC, we did not detect this species north of La Paz (24.17° N).

***Echinolittorina penicillata***

Historically, the northern range limit of *Echinolittorina penicillata* the EP coast was located at Bahía Magdalena (24.62° N), with the distribution extending southwards to the Jalisco region of Mexico (Reid, 2002). This species was also relatively common throughout the GoC (Reid, 2002). Our surveys revealed that *E. penicillata* remains common throughout the GoC, and its modern, northern range limit now extends up to Punta Morro Santo Domingo (28.24 ° N) on the EP coast, suggesting a recent range expansion of ~816 km.

***Epitonium acapulcanum***

The historical geographic range of *Epitonium acapulcanum* extends from Bahía Magdalena (24.62° N), into the GoC, and south to the Galapagos Island (Keen 1971). Throughout our surveys, we have only identified a single individual across all of the visited sites in Baja California (Punta Marquez; 23.96 ° N). However, as this is a largely understudied species (only three, relatively conspicuous occurrences exist in the GBIF database for Baja California; GBIF.org, 2025a), it not possible to reliably infer any potential changes in distributions with a limited species record (both historically, and in modern surveys).

***Eupleura muriciformis***

According to Keen (1971), historically, this species was present throughout the GoC, extending its distribution southward towards Ecuador. During our exhaustive surveys, we have only identified individuals at La Gringa (29.05 ° N) within the GoC. However, this species occupies both intertidal rocky shores and offshore reefs (Keen, 1971), and consequently, our intertidal surveys could not capture its true distribution.

***Fissurella rubropicta***

Historically, the distribution of *Fissurella rubropicta* extended from its northern range limit at Lagoon Head, Baja California (~ 28° N), south to Oaxaca, Mexico (~ 16° N) (Keen, 1971). This species has demonstrated a northwards range expansion since, with its northern limit now located between Santa Rosalillita and Punta Baja (28.6–29.6° N; Zarzyczny et al., 2024).

***Fissurella volcano***

Historical distribution of this species ranged from Crescent City, California (41.8° N; Morris et al., 1980), south to Punta Marquez (24.0° N) (Groves and Mertz, 2025). Since, this keyhole limpet has retracted its range by ~144 km within Baja California, to Bahía Magdalena (24.6° N; Zarzyczny et al., 2024).

***Haliotis fulgens***

Historically, the distribution of *Halotis fulgens* ranges from Point Conception (34.45 ° N) to Bahía Magdalena (24.62° N; Morris, 1980). We detected individuals at Bahía Magdalena and San Juanico only, however it is worth noting that this species occupies the low intertidal zone, and the shallow subtidal down to 10m depth (Morris, 1980). Nevertheless, our surveys suggest no range retraction at the southern range limit has occurred.

***Heliacus areola***

Historically, the distribution of *Heliacus areola* ranges from La Paz (24.17° N) within the GoC, extends north throughout the GoC, and then south along the coast of Mexico to Panama and Galapagos Islands (Keen, 1971). This species is absent from the EP coast (Keen, 1971). We only detected this species at Agua Verde (25.52° N). However, as *H. areola* occupies the extreme low tide zone only (Keen, 1971), our ability to capture the true distribution of this species was likely limited.

***Jenneria pustulata***

The historical distribution of this distinctive cowry extends from the northern portion of the GoC, and south to Ecuador (Keen, 1971). We detected *Jenneria pustulata* on the EP coast, at Punta Marquez (23.96 ° N), with further confirmed records of this species in this area from in 2023 (iNaturalist, 2025a). The repeated detection of this snail (albeit in small numbers) could be indicative of a recent range expansion from the GoC, to the EP coast.

***Liocerithium judithae***

Historically, Keen (1971), describes *Liocerithium judithae* as present at Bahía Magdalena (24.6° N) on the EP coast, with a distribution through the GoC, and south to the Matzalan region of Mexico. There are several collections of *L. judithae* catalogued at the LACM (e.g. LACM 1967-72.15, 1967-73.10, and LACM 1966-8.18; Groves & Mertz, 2025). However, whilst we detected this species across all our surveyed sites within the GoC, we did not detect it on the EP coast. There are also no modern observations of this species on the EP coast on iNaturalist (iNaturalist, 2025b). Our findings therefore point towards a potential local extinction of this species at Bahía Magdalena.

***Littorina keenae***

According to Lee and Boulding (2007), the historic distribution of *Littorina keenae* ranged from Charleston, Oregon (43.3° N) to at least Laguna Manuela (28.2° N), with occasional records further south as far as north side of Bahía Magdalena (~24.8° N). However, we have identified further two records of *L. keenae* from SBMNH (catalogue numbers 369746 and 369754; Chaney et al., 2024), collected from Cabo San Lucas (22.95° N). During our surveys, we consistently detected *L. keenae* between (and including) Punta Abreojos (26.71° N), and Campo Kennedy (31.71° N). Whilst we did not detect any individuals at Bahía Magdalena (24.62° N), we have identified occasional individuals as far south as Pozo de Cota (23.02° N). Our findings therefore suggest that the distribution of this species remains largely unchanged.

***Littorina scutulata***

Historically, the distribution of *Littorina scutulata* extended from Alaska (Lopez, 2025) as far south as San Juanico (26.23° N; Groves and Mertz, 2025; LACM 1971-6.17). We consistently detected *L. scutulata* in the northern portion of the Baja California Peninsula, and established Bahía Asunción (27.13° N) as the present southern range limit, suggesting a range retraction of ~ 240 km.

***Lottia acutapex***

Historically, this limpet is present throughout the GoC (Keen, 1971). We detected *Lottia acutapex* across all surveyed sites within the GoC, with no observations on the EP coast, suggesting the distribution of this species remains unchanged. Records of this species on the EP coast, are most likely *Lottia conus*, which shares morphological similarities with this species.

***Lottia argrantesta***

When Simison and Lindberg (2003), first described *Lottia argrantesta,* they described the distribution of this species as restricted to the GoC, with records from La Paz (~24° N), north to San Francisquito (~28° N), and back south to Guayamas (~27° N) on the Sonoran coast. We have consistently detected this limpet across all three surveyed sites within the Gulf of California, suggesting the distribution of this species since its first description remains largely unchanged. Although we acknowledge that given our sparse survey locations within the GoC, we are unlikely to determine any small scale range changes limited to the GoC.

***Lottia asmi***

The historical distribution of this temperate limpet extends from Canada (Groves and Mertz, 2025; LACM 1968-67.2) as far south as San Juanico (26.2° N) with records of this species from 1979 (Groves and Mertz, 2003; LACM 1979-25.11) and 1971 (Groves and Mertz 2023; LACM 1971.6.10). Whilst we regularly detected *Lottia* *asmi* on rocky shores along the EP coast, we did not detect this species south of Punta Abreojos (26.71° N), suggesting a range retraction of ~ 120 km.

***Lottia atrata***

According to Keen (1971), the historical distribution of this species along the Baja California coastline is discontinuous. At the time of the publication, Keen (1971) describes *Lottia atrata* as common at Bahía Magdalena (24.6° N) on the EP coast, and in the northern portion of the GoC (31.5° N), and stretching south along the east coast of the GoC to Kino Bay, Sonora (28.9° N). Whilst we did not survey the east coast of the GoC, we did detect *L. atrata* on the west coast at La Paz (24.17° N) and Agua Verde (25.52° N). On the EP coast, we detected *L. atrata* at Bahía Magdalena (24.6° N), and Cerritos Point (23.33° N). Our findings suggest a potential new population at Cerritos Point, but not a poleward range expansion.

***Lottia austrodigitalis***

According to Kelly and Palumbi (2010), the southern limit of *Lottia austrodigitalis* is between between Cerritos Point (23.33 ° N) and Pozo de Cota (23.02 ° N) at ~23.2° N. Although the work was published in 2010, the reported species ranges are based on primary literature (Kelly and Palumbi, 2010), and likely reflect historical distribution. In our field surveys, *L. austrodigitalis* was common along the EP coast. However, we did not detect this limpet south of Punta Abreojos (26.71° N), suggesting a substantial range retraction of ~ 648 km.

***Lottia conus***

The historical records of this species are scarce, meaning that reliable historical range limits cannot be determined (Zarzyczny et al., 2024). Consequently, we provide the modern distribution of this species as reported by Zarzyczny et al., (2024). *Lottia conus* has a relatively continuous distribution across the rocky shores on the EP coast, from its southern range limit at Bahía Magdalena (24.62 ° N), north to at least Jalama Beach, CA (34.5° N).

***Lottia dalliana***

According to Keen (1971), the historic distribution of *Lottia dalliana* was limited to the northwestern portion of the GoC, from Puerto Peñasco (31.3° N) in the northern Gulf, southwest to San Francisquito Bay (28.4° N). Whilst we did not survey the GoC extensively, we did record this species at La Gringa (29.05° N) only.

***Lottia fascicularis***

We detected *Lottia* *fascicularis* on the EP coast between Bahía Magdalena (24.6° N) and Pozo de Cota (23.02 ° N; southernmost surveyed site), although the distribution of this species extends south to Costa Rica (Keen, 1971). Historically, the northernmost record of this limpet is also from Bahía Magdalena (24.6° N; ANSP catalogue no. 152206; Rosenberg, 2025) suggesting the distribution of this species (at least in the northern portion of its range) remains unchanged.

***Lottia fenestrata***

The range of this temperate limpet extends from the EP coast of USA, south into Baja California. Historically, the southernmost record of *Lottia fenestrata* was collected in 1954 (LACM 1954-14.8; Groves and Mertz, 2025) at Bahía San Bartolome (27.66° N) near Punta Eugenia. However, in our surveys we have consistently detected this species south of this locality at Bahía Asunción (27.13° N), Punta Abreojos (26.71° N), San Juanico (26.24° N) and La Bocana (26.06° N). It is worth acknowledging that historical records of this species are sparse within Baja California (a total of nine records exist within the GBIF database; GBIF 2025b). Moreover, given cryptic diversity exists within the *Lottia* genus, smaller individuals of this species can be confused with *Lottia strigatella* which also often exhibit an eroded apex. Whilst we cannot exclude a southward range expansion of this species, given the limited historic records combined with cryptic diversity of the genus, we cannot reliably determine a historic distribution of *Lottia fenestrata.*

***Lottia gigantea***

Please see the main text for an in depth overview of this species.

***Lottia limatula***

The range of this temperate species extends from Canada, south to Baja California Sur. The southernmost historical records were collected in 1979 (LACM 1979-25.14; Groves and Mertz 2023) and 1971 (LACM 1971-6.11; Groves and Mertz 2023) at San Juanico (26.23° N). Whist the northern range limit of this species falls outside of our geographic study area, in the southern portion of this species’ range, we have consistently detected *Lottia limatula* between UABC (31.86° N) and Punta Abreojos (26.71° N). We did not detect *L. limatula* at San Juanico (26.24° N) or any other southern sites. We can therefore infer a range contraction of 120 km from San Juanico to Punta Abreojos.

***Lottia mesoleuca***

The range of this tropical limpet extends from Galapagos Islands, north along the EP coastline to Matzalan, Mexico and into the GoC at Cerralavo Island, and tip of the Baja California Peninsula (Keen, 1971). Historically, the northern range limit of *Lottia mesoleuca* on the coast of Baja California Peninsula was at Cabo San Lucas (22.9° N) at the tip of the Peninsula (Keen, 1971). We detected *Lottia mesoleuca* on the EP coast of Baja California at Pozo de Cota (23.02° N), suggesting a potential range expansion of ~48 km from Cabo San Lucas.

***Lottia pediculus***

According to Keen (1971), the historic distribution of this species ranges from Espiritu Santo Island (~24.5° N) in the GoC, south to Colombia. Whilst we did not survey the Espiritu Santo Island, we did detect *Lottia pediculus* at La Gringa (29.05° N) suggesting potentially a substantial range expansion within the GoC of ~960 km.

***Lottia pelta***

According to Morris (1980), the distribution of *Lottia pelta* ranges from the Aleutian Archipelago, south to Bahía del Rosario, just south of Punta Baja (29.95° N). We consistently detected *L. pelta* between UABC (31.86° N) to Punta Baja (29.95° N), suggesting that the range of this species remains unchanged.

***Lottia scabra***

This temperate limpet shares cryptic morphology with *Lottia conus*, albeit it exhibits a more northern distribution extending from California, south into Baja California (Coombs, 2017). Southernmost historical record (ANSP Catalogue Number: 39105; Rosenberg, 2025) of this species is from Punta Abreojos (26.71° N) albeit the collection date of the specimen is unknown. In our surveys, we established the southern limit of this species at Punta Caballo de Piedra (27.81° N), with no records of this limpet on any sites further south, including Punta Abreojos. It is possible that the southern records of this species could be confused with *Lottia conus*; for example, some older publications suggest the range of *Lottia scabra* extends south to Cabo San Lucas (22.88° N; Shanks et al., 2014; Morris et al., 1980) which is most likely based on historical misidentification of *Lottia conus.* However, if we assume the historical records from Punta Abreojos are correct, our results suggest a range retraction of ~336 km has occurred from Punta Abreojos to Punta Caballo de Piedra.

***Lottia stanfordiana***

According to Keen (1971), the distribution of *Lottia stanfordiana* ranges from the head of the GoC, south to Guayamas and Isla Espíritu Santo. Our surveys detected this limpet at La Gringa (29.05° N) only. With no records in the southern Gulf sites such as Agua Verde (25.52° N) or La Paz (24.17° N). Although we surveyed the GoC sparsely, the absence of *L. stanfordiana* from southern GoC sites could be indicative of changes in distribution of this species.

***Lottia strigatella***

The historical distribution of *Lottia strigatella* is unclear due to the absence of reliable historical records associated with cryptic species diversity (Zarzyczny et al., 2024). Zarzyczny et al., 2024, placed the current northern range limit of *L. strigatella* as somewhere between San Diego (32.70° N) and Punta Baja (29.95° N), with the distribution extending south the Pozo de Cota (23.02° N). However, in our recent surveys, we obtained samples of *L. strigatella* from northern Baja California, including from Campo Kennedy (31.70° N), and Universidad Autónoma de Baja California, Ensenada (31.86° N). We can therefore refine the northern range limit of *L. strigatella* as somewhere between Ensenada (31.86° N) and San Diego (32.70° N).

***Lottia strongiana***

According to Keen (1971), the distribution of this species is limited to the GoC with the range extending from the head of the Gulf, south to Puerto Libertad (29.9° N) on the east coast, and Cabo San Lucas (22.88° N) on the west of the Goc. We only detected this species at La Gringa (29.05° N), and not at Agua Verde (25.52° N) or La Paz (24.17° N), however, given that we surveyed a limited number of sites within the GoC, it is difficult to reliably determine whether a range change has occurred.

***Lottia turveri***

Historically, the distribution of *Lottia turveri* was limited to the GoC, with the range extending from the head of the Gulf, south to Guayamas (27.9° N) in the east, and Bahía de los Angeles (28.95° N) in the west. We detected this limpet across all of the surveyed sites in the Gulf (La Gringa, 29.05° N; Agua Verde, 25.52° N; and La Paz, 24.17° N) as well as two further sites on the EP coast (Cerritos Point, 23.33° N; and Punta Marquez 23.96° N). The new records suggest a substantial range expansion, both within the Gulf (southward), and across to the EP (poleward), totalling to ~ 240 km range expansion.

***Macron aethiops***

According to Keen (1971), *Macron aethiops* is common on the EP coast of Baja California, and rare in the GoC. Based on the GBIF collection (GBIF.org, 2025c), the historical distribution of *M. aethiops* spans from Bahía San Quintin (30.48° N; SBMN 626455, SBMN 366611 and LACM 1973-2.25, collected in 1950, 1971 and 1973, respectively) to Cabo San Lucas (22.88° N; SBMNH Catalogue number 146456; collected 1982) on the EP coast. Within the GoC, occasional records exist from La Paz (24.19° N; MCZ 23748, no date) and Isla Danzante (25.79° N; SBMNH 152682, collected in 1978), albeit the latter is a dry specimen, collected on SCUBA. During our surveys, we detected *M. aethiops* only at Punta Abreojos (26.71° N) only which can likely be attributed to the habitat preference of this species. Whilst *M. aethiops* does reside on rocky shores, the this whelk often inhabits habitats with softer sediments such as mudflats (Rodriguez, 2009), which we did not capture in our surveys.

***Macron lividus***

Historically, the range of *Macron lividus* extended from Newport, California (39.58° N; MCZ Catalogue Numbers 132122 and 327488), south to San Juanico (26.25° N; LACM 1955-5.9, LACM 1971-181.18, LACM 1979-25.24, MCZ Catalogue Numbers 327490 and 327491;GBIF.org, 2025d). In our field surveys, we did not detect *M. lividus* at San Juanico or nearby La Bocana (26.06° N). Our southernmost record of this whelk is Punta Abreojos (26.71° N), suggesting a range retraction of ~120 km has occurred at the southern range limit of this species.

***Megaastrea undosa***

Historical distribution of *Megaastrea unodosa,* extends from southern California where it is relatively common, south along the EP coast to Punta Abreojos (Keen, 1971). We detected this species across the coast of Baja California Norte, and as far south as Punta Abreojos (26.71° N) suggesting the range of *M. undosa* within Baja California remains unchanged.

***Megathura crenulata***

The reported range of this species is Point Conception, California (34.35° N), south to Isla Asunción, Baja California (27.10° N; Bonett-Calzada et al., 2024). We detected this species at Campo Kennedy (31.70° N) and La Chorera (30.47° N) only. However, it is worth acknowledging that this species occupies both the intertidal and shallow subtidal regions (Gotshall, 2005), therefore our surveys were unlikely to capture the full extent of this species’ distribution.

***Mexacanthina angelica***

Historically distributed throughout the GoC (Keen, 1971), from the head of the Gulf, south to Puerto Escondido (25.7° N; LACM 1937-71.5; Groves and Mertz, 2025). We detected *Mexacanthina angelica* at La Gringa (29.05° N) only. However, the two other GoC sites which we surveyed, Agua Verde (25.52° N) and La Paz (24.17° N) fall outside of the historic range of this species.

***Mexacanthina lugubris***

A change in the distribution of *Mexacanthina lugubris* has already been reported by Fenberg et al. (2014). Historically, the range of this whelk extended from southern California to Bahía Magdalena (Keen, 1971). Fenberg et al. (2014) reported a range retraction of ~ 331 km from Bahía Magdalena (24.6° N) to La Bocana (26.06° N). As we did not identify any individuals of *M. lugubris* at La Bocana, our survey results indicate a further 33 km range retraction to San Juanico (26.24° N).

***Mitrella delicata***

According to Keen (1971), *Mitrella delicata* was historically distributed from Guayamas, Mexico (~27.9° N), along the Mexican coastline and south to Panama. This species was absent from Baja California, with the exception of a single historic record from Caleta San Lucas (27.20° N; SBMNH catalogue number 154785; Chaney et al. 2024) in the GoC. We have identified this species on the EP coast of Baja California, at Cerritos Point (23.33° N), Punta Marquez (23.96° N), and as far north as La Bocana (26.06° N). Whilst we did not observe *M. delicata* at San Juanico (26.24° N), verified observations of this species exist from the vicinity of this location (Simison, 2018). Our surveys, combined with verified observations of this species suggest a substantial range expansion of at least 648 km.

***Mitrella ocellata***

*Mitrella ocellata* has a wide distribution, spanning both EP and Atlantic coasts (GBIF.org, 2025e). In the EP, the historic distribution of this species extends from the Galápagos (-0.40° N), north along the coast of Mexico, into the GoC where it is found along the east and west coasts and the head of the Gulf (GBIF.org, 2025e). On the EP coast of the Baja California Peninsula, the historic range limit of *M. ocellata* is at Cabo San Lucas (22.87° N; INHS Catalogue number 56298; GBIF.org, 2025e). During our surveys, we detected this species as far north on the EP coast as Cerritos Point (23.33° N), suggesting a range expansion of at least ~72 km

***Modulus cerodes***

*Modulus cerodes* is historically reported as present on mudflats throughout the GoC, south to Panama (Keen, 1971). We only detected an individual on the coast of the GoC, at Agua Verde (24.52° N). However, given our surveys were limited to rocky shores, we did not capture the true distribution of this species.

***Neorapana tuberculata***

According to Keen (1971), historically, this species is restricted to the GoC, from Cabo San Lucas to Mazatlán. We only detected a single individual at La Gringa (29.05° N) in the GoC. However, as *Neorapana tuberculata* is often found subtidally, our intertidal surveys did not capture the true distribution of this species.

***Neoterebra variegata***

*Neoterebra variegata* has a broad Panamic distribution, extending from GoC (GBIF.org, 2025f), south to Panama (Keen, 1971). This auger snail inhabits both the intertidal and subtidal zone (GBIF.org, 2025f). During our surveys, we detected this snail at Agua Verde (25.52° N) only. However, as our surveys focussed on the intertidal zone, we did not capture the true distribution of this species in our study.

***Nerita funiculata***

Historically, the distribution of *Nerita funiculata* ranged from Peru (18.4° S–0°; Keen, 1971), north to Punta Morro Santo Domingo on the EP coast (26.30° N; Groves and Mertz, 2023f), with a vast distribution throughout the GoC (Keen, 1971). As reported by Zarzyczny et al., (2024), the northern range limit of this species has recently expanded on the EP coast by 432 km, to Punta Caballo de Piedra (27.8° N).

***Nerita scabricosta***

Historically, the distribution of *Nerita scabricosta* ranged from Ecuador (3.0° S–1.0° N; Keen, 1971), north to Punta San Hipolito on the EP coast (27.0° N; Groves and Mertz, 2025), with a vast distribution throughout the GoC (Keen, 1971). As reported by Zarzyczny et al., (2024), the northern range limit of this species has recently expanded on the EP coast by 240 km, to Punta Caballo de Piedra (27.8° N).

***Norrisia norrisii***

Historical literature suggests the range of *Norrisia norrisii* extends from Point Conception, California (34.4° N), south to Isla Asunción, Baja California (27.1 ° N; Stebbins, 1986). *Norrisia norrisii* is a subtidal herbivore, which occasionally occupies the lower intertidal zone (Hartman, 1983). We identified a single individual at Punta Baja (29.95 ° N), however, given that our surveys were limited to the rocky intertidal, we are unable to capture the true modern distribution of this species.

***Notocochlis chemnitzii***

Historically, the distribution of *Notocochlis chemnitzii* is reported as ranging from Bahía Magdalena (24.6° N), south down the EP coastline, through the GoC and south to Galapagos Islands (Keen, 1971). We only detected a single individual at Agua Verde (25.52° N) within the GoC, however, as this moon snail inhabits intertidal muddy sediments (Keen, 1958; Smith and Dietl, 2015), our surveys did not capture its true modern distribution.

***Olivella intorta***

According to Keen (1971), the historical distribution of *Olivella intorta* extends from Bahía Magdalena (24.62° N) south to the tip of Baja California and into the southern end of the GoC. Occurrences along west coast of the GoC exists do not extend north of Cabo Pulmo (23.44° N; UF 557220-Mollusca, Florida Museum of Natural History, 2025; ANSP 477543, Rosenberg, 2025). On the east side of the GoC, occurrences of *O. intorta* are limited to Mazatlan (23° N; SDNHM 43815; San Diego Natural History Museum, 2025), and Isla de la Raza (27.9° N; BMSM 61433; BMSM, 2025). We detected *O. intorta* at Agua Verde (25.52° N) only, however, given this genus prefers sandy or muddy substrate (Kelly et al., 2021; Corte et al., 2020) it is likely that our rocky shore surveys did not capture the true modern distribution. Nevertheless, our detection of this gastropod at Agua Verde may be indicative of a northwards range expansion of at least ~ 456 km on the west coast of the GoC.

***Parvanachis diminuta***

Historically distributed on the EP coast from Punta Abreojos (26.71° N) to Panama (9° N); Keen, 1971). We detected a small population at San Juanico (26.24 ° N) only. However, as this small species typically occupies muddy sediment (Maintenon, 2014), our rocky shore surveys likely did not capture its true distribution.

***Penethais triangularis***

Historically, *Penethais triangularis* was distributed from Cabo San Lucas (22.88° N) on the EP coast, around the tip of the Peninsula, through the GoC and south to Peru. We detected this species at Cerritos Point (23.32 ° N) suggesting a range expansion of ~96 km along the EP coastline. Interestingly, fossils dating back to the Pleistocene have been found at Bahía Magdalena (24.62° N; Roopnarine and Garcia, 2024) suggesting that this species has undergone a range contraction since, likely associated with the Pleistocene climatic fluctuations (Hofreiter and Stewart, 2009), and is now once again expanding polewards with the ongoing tropicalisation of the peninsula.

***Plicopurpura columellaris***

Keen (1971) reports the historic northern range limit as Bahía Magdalena (24.62° N), however specimens collected in 1979 (LACM 1979-25.13; LACM 1979-24.8) from the Punta Pequeña region (26.2 - 26.3° N) are preserved at the LACM (Groves and Mertz, 2025). We detected populations of *Plicopurpura columellaris* north of its historic range limit, as far as Punta Caballo de Piedro (27.81° N), suggesting a range expansion of ~456 km.

***Polinices uber***

*Polinices uber* occurs both intertidally and offshore (up to 90m), throughout the GoC and along the EP coastline of Baja California (Keen, 1971). Consequently, whilst we detected this species in the GoC at Agua Verde (25.52° N), our surveys could not capture its true modern distribution.

***Pomaulax gibberosus***

According to Keen (1971), the historic range of *Pomaulax gibberosus* extends from Queen Charlotte Islands, British Columbia (52-54° N), south to Bahía Magdalena (24.6° N). Whilst *P. gibberosus* occurs intertidally in the northern portion of its range, the southern populations can be subtidal, at depths greater than 17m (Keen, 1971). During our surveys, we found an individual intertidally at Punta Abreojos (26.71° N) only. However, given the subtidal distribution of this species in Baja California Sur, our surveys could not capture this snail’s true modern distribution.

***Pusio elegans***

According to Keen (1971) historic distribution of this species ranges from Bahía Magdalena (24.62° N), throughout the GoC and south to Peru. Whilst we did not discover any individuals at our three surveyed sites in the GoC, we did detect *Pusio elegans* north of Bahía Magdalena, at Las Barrancas (26.0° N), and San Juanico (26.23° N), suggesting a recent northward range expansion of 264 km.

***Rissoina stricta***

The distribution of this species ranges from Cabo San Lucas (22.88° N) at the tip of Baja Peninsula, and throughout the GoC, south to Galapagos Islands (Galapagos Species Database, 2025; Keen, 1971). We detected a single individual within the GoC at Agua Verde (25.52° N). However, as *Rossoina stricta* occurs subtidally, up to 200m (Galapagos Species Database, 2025) our surveys could not capture its true distribution.

***Roperia poulsoni***

The historic distribution of *Roperia poulsoni*, extends from California, south to Bahía Magdalena (24.63° N; SIO catalogue number M4131; Seid, 2025). We did not detect any individuals south of San Juanico (26.23° N), in any of our exhaustive surveys or in the iNaturalist database, suggesting a range retraction of ~264 km.

***Siphonaria maura***

The distribution of *Siphonaria maura* extends from Peru, north along the EP of Mexico, throughout the GoC, and to the historic northern range limit at Bahía Magdalena (24.63° N; ANSP catalogue number 151709; Rosenberg, 2025). In our surveys, we detected *S. maura* throughout the GoC, and along the EP coast of Baja Califronia. On the EP coast, we detected *S. maura* beyond its northern historic range limit, at San Juanico (26.23° N) and Punta Conejo (24.05 ° N), suggesting a range expansion of ~264 km by this species.

***Stramonita biserialis***

Historical northern range limit for this species is Punta Eugenia (27.9° N; Keen, 1971; Fenberg et al., 2023). We consistently detected populations of *Stramonita biserialis,* north of the historical range limit, and as far Punta Baja (29.95° N). However, the modern northern range limit of this species is thought to extend to Orange County, CA based on recent confirmed records (33.5° N: inaturalist.org/observations/9674212), as reported by Fenberg et al., (2023). This observation suggests a potential substantial range expansion of ~1008 km.

***Strigatella tristis***

According to Keen (1971), the historic distribution of this species ranges from the northern end of the GoC south to Ecuador. However, several dry specimens have been collected by C.R. Orcutt from Bahía Magdalena (24.6° N) in 1917 which are catalogued at the ANSP (Catalogue number 151749; Rosenberg, 2025) identity of which we were able to confirm as *Strigatella tristis.* Moreover, fossil records of *S. tristis* from Bahía Magdalena, dating back to the Pleistocene exist (CASG:FOSSIL:859-60; Roopnarine and Garcia, 2024). Our surveys detected *S. tristis* both on the EP coast at Punta Marquez (23.95 ° N) and in the GoC at La Gringa (29.05 ° N). However, given the uncertain and patchy historic distribution of this species, we are unable to make reliable inferences of any potential range changes.

***Tegula aureotincta***

The historic range of *Tegula aureotincta* extends from Bahía Magdalena (24.62 ° N), north to Ventura County, California (~34 ° N; Hellberg, 1998). In our surveys, we consistently detected *T. aureotincta* between Bahía Magdalena and our northernmost surveyed site, UABC (31.86° N). The distribution of this species along the EP coast of Baja California Peninsula remains unchanged.

***Tegula funebralis***

Historic range of *Tegula funebralis* extends from Vancouver to Central Baja on the EP coast (Hellberg, 1998). The southernmost record of *T. funebralis* species is from San Juanico (26.25 ° N; SBMNH Catalogue Number 158841; Chaney et al., 2024) where we also detected it in our surveys, suggesting the distribution of this species remains unchanged.

***Tegula gallina***

As reported by Zarzyczny et al. (2024), *Tegula gallina* has experienced a range contraction along the EP coast of Baja California. The historic range of this species extended from Point Conception, CA (34.4° N) to Punta Marquez (24.0° N) (Keen, 1971; Zarzyczny et al., 2024). The modern southern range limit for *T. gallina* is currently Bahía Magdalena (24.6° N), demonstrating a range expansion of ~144 km (Zarzyczny et al., 2024).

***Tegula rugosa***

Historically, the distribution of *Tegula rugosa* extends from the head of the GoC, south to Bahía Conception (27 ° N). We detected *T. rugosa* at La Gringa (29.05 ° N), however, as the next southernmost site which we surveyed (Agua Verde; 25.52 ° N) falls beyond the historic southern range limit of this species, we are unable to establish whether the range of this species has changed.

***Trachypollia lugubris***

*Trachypolia lugubris* has a broad distribution, historically extending from San Diego, California, south to Panama along the EP coast (Keen, 1971). We only detected this species at Santa Rosalillita (28.65 ° N), however, as *T. lugubris* occurs at depths of up to 40m (Keen, 1971) we are unable to determine the true modern range of this species from our intertidal surveys.

***Turbo fluctuosus***

According to Keen (1971), the historic range of *Turbo fluctuosus* is discontinuous, stretching south from Isla de Cedros (28 ° N), on the EP coast of Baja California, throughout the GoC, and south to Peru. We identified individuals throughout the Gulf of California, at La Gringa (29.05 ° N) and La Paz (24.20 ° N), as well as along the EP coast from Punta Marquez (23.96 ° N) to San Juanico (26.23° N) suggesting the distribution remains discontinuous albeit unchanged.

***Turbo funiculosus***

Historically, *Turbo funiculosus* has not been observed in Baja California as frequently as *T. fluctuosus* (Keen, 1971). The historic range limit of this species appears to be located as Punta Pequena (26.23° N) on the EP coast of Baja California (LACM 1979-25.9; Groves and Mertz, 2025). We have detected *T. funiculosus* north of the historic range limit at Punta Abreojos (26.71° N) and Punta Caballo de Piedro (27.81 ° N) suggesting a recent range expansion of ~456 km.

***Uvanilla olivacea***

Historically, the distribution of *Uvanilla olivacea* ranges from Cabo San Lucas (22.88 ° N) on the southern tip of the Baja California Peninsula, to La Paz (24.20 ° N) on the east coast of GoC, and Matzalan on the west coast of the Gulf, extending south to Oaxaca (Keen, 1971). As this species inhabits both the rocky intertidal and shallow subtidal habitats, our surveys were unable to determine the full extent of its modern distribution in Baja California. However, we detected *U. olivacea* at Punta Marquez (23.96 ° N), suggesting a range expansion of at least 192 km.

***Vasula speciosa***

The historic distribution of *Vasula speciosa* ranges from Bahía Magdalena (24.62 ° N), south along the EP coast, throughout the GoC and south to Peru. We detected *V. speciosa* both within the GoC at La Gringa (29.05 ° N), and along the EP coastline, as far north as San Juanico (26.24 ° N) suggesting a range expansion of 264 km.

***Vitta luteofasciata***

*Vitta luteofasciata* typically inhabits mudflats and mangrove swamps along the GoC, south to Peru (Keen 1971). Whilst our surveys of rocky shores were unable to capture the true distribution of this species, we did record *V. luteofasciata* at La Paz within the GoC (24.20 ° N).

***Zetecopsis zeteki***

The historic distribution of *Zetecopsis zeteki* ranges from the head of the GoC south to Ecuador (Keen, 1971). We detected this species at Agua Verde (25.52 ° N) only. However, as this species can be present subtidally (GBIF.org, 2025h), our surveys did not capture the full distribution of this species within the Baja California region.

**Supplementary Tables**

**Table S1.** The surveyed sites, site coordinates and the years when the exhaustive surveys took place.

| **Site** | **Site Code** | **Latitude** | **Longitude** | **Years Surveyed** |
| --- | --- | --- | --- | --- |
| **Eastern Pacific Coast** | |  |  |  |
| U.A.B.C | UA | 31.86167 | 116.6681 | 2024 |
| Campo Kennedy | CK | 31.70278 | -116.684 | 2024 |
| La Chorera | LCH | 30.4700 | -116.046 | 2024 |
| Punta Baja | PB | 29.9516 | -115.812 | 2021-2022, 2024 |
| Santa Rosalillita | SR | 28.6533 | -114.246 | 2021-2022, 2018 |
| Punta Morro Santo Domingo | PMSD | 28.2421 | -114.094 | 2018 |
| Punta Caballo de Piedro | PCP | 27.8119 | -114.774 | 2017 |
| Punta Clambey | PCL | 27.6510 | -114.845 | 2017 |
| Bahía Asunción | BA | 27.1337 | -114.305 | 2021-2022, 2018 |
| Punta Abreojos | PA | 26.7104 | -113.573 | 2021-2022, 2017 |
| San Juanico | SJ | 26.2377 | -112.479 | 2017, 2018, 2024 |
| La Bocana | LB | 26.0571 | -112.286 | 2018 |
| Las Barrancas | LBR | 25.9993 | -112.204 | 2021-2022 |
| Bahía Magdalena | BM | 24.6181 | -112.134 | 2021-2022 |
| Punta Conejo | PC | 24.0546 | -110.982 | 2018 |
| Punta Marquez | PMZ | 23.9554 | -110.873 | 2021-2022, 2017 |
| Cerritos Point | CP | 23.3292 | -110.179 | 2021-2022, 2017 |
| Pozo de Cota | PDC | 23.0194 | -110.098 | 2021-2022, 2017, 2024 |
| **Gulf of California** | |  |  |  |
| La Paz | LP | 24.1703 | -110.31 | 2021-2022 |
| Agua Verde | AV | 25.5243 | -111.102 | 2021-2022, 2017 |
| La Gringa | LG | 29.0512 | -113.534 | 2021-2022, 2018 |

**Table S2.** A list of species detected during the exhaustive surveys of the rocky intertidal along the coast of Baja California Peninsula. The surveys took place between 2017 and 2024. Species are listed the alphabetical order based on family.

| Family | Subfamily | Species |
| --- | --- | --- |
| Architectonicidae |  | *Heliacus areola* |
| Buccinoidea |  | *Macron aethiops* |
| Buccinoidea |  | *Macron lividus* |
| Cerithiidae | Cerithiinae | *Cerithium adustum* |
| Cerithiidae | Cerithiinae | *Cerithium maculosum* |
| Cerithiidae | Cerithiinae | *Cerithium menkei* |
| Cerithiidae | Cerithiinae | *Cerithium stercusmuscarum* |
| Cerithiidae | Cerithiinae | *Liocerithium judithae* |
| Columbellidae |  | *Anachis berryi* |
| Columbellidae |  | *Anachis vexillum* |
| Columbellidae |  | *Costoanachis sp.* |
| Columbellidae |  | *Columbella aureomexicana* |
| Columbellidae |  | *Columbella fuscata* |
| Columbellidae |  | *Columbella major* |
| Columbellidae |  | *Columbella socorroensis* |
| Columbellidae |  | *Columbella strombiformis* |
| Columbellidae |  | *Mitrella delicata* |
| Columbellidae |  | *Mitrella ocellata* |
| Columbellidae |  | *Mitrella sp.1* |
| Columbellidae |  | *Parvanachis diminuta* |
| Conidae |  | *Conus californicus* |
| Conidae |  | *Conus nux* |
| Epitoniidae |  | *Epitonium acapulcanum* |
| Fissurellidae | Fissurellideinae | *Diodora digueti* |
| Fissurellidae | Fissurellideinae | *Fissurella rubropicta* |
| Fissurellidae | Fissurellideinae | *Fissurella sp.* |
| Fissurellidae | Fissurellideinae | *Fissurella volcano* |
| Fissurellidae | Diodorinae | *Megathura crenulata* |
| Haliotidae |  | *Haliotis fulgens* |
| Hipponicidae |  | *Antisabia panamensis* |
| Littorinidae | Littorininae | *Echinolittorina albicarinata* |
| Littorinidae | Littorininae | *Echinolittorina apicina* group |
| Littorinidae | Littorininae | *Echinolittorina aspera* group |
| Littorinidae | Littorininae | *Echinolittorina modesta* |
| Littorinidae | Littorininae | *Echinolittorina penicillata* |
| Littorinidae | Littorininae | *Littorina keenae* |
| Littorinidae | Littorininae | *Littorina scutulata* |
| Lottiidae | Lottiinae | *Lottia acutapex* |
| Lottiidae | Lottiinae | *Lottia argrantesta* |
| Lottiidae | Lottiinae | *Lottia asmi* |
| Lottiidae | Lottiinae | *Lottia atrata* |
| Lottiidae | Lottiinae | *Lottia austrodigitalis* |
| Lottiidae | Lottiinae | *Lottia conus* |
| Lottiidae | Lottiinae | *Lottia dalliana* |
| Lottiidae | Lottiinae | *Lottia fascicularis* |
| Lottiidae | Lottiinae | *Lottia fenestrata* |
| Lottiidae | Lottiinae | *Lottia gigantea* |
| Lottiidae | Lottiinae | *Lottia limatula* |
| Lottiidae | Lottiinae | *Lottia mesoleuca* |
| Lottiidae | Lottiinae | *Lottia pediculus* |
| Lottiidae | Lottiinae | *Lottia pelta* |
| Lottiidae | Lottiinae | *Lottia scabra* |
| Lottiidae | Lottiinae | *Lottia stanfordiana* |
| Lottiidae | Lottiinae | *Lottia strigatella* |
| Lottiidae | Lottiinae | *Lottia strongiana* |
| Lottiidae | Lottiinae | *Lottia turveri* |
| Mitridae | Strigatellinae | *Strigatella tristis* |
| Modulidae |  | *Modulus cerodes* |
| Muricidae | Ocenebrinae | *Acanthinucella punctulata* |
| Muricidae | Ocenebrinae | *Acanthinucella spirata* |
| Muricidae | Ergalataxinae | *Claremontiella nodulosa* |
| Muricidae | Coralliophilinae | *Coralliophila nux* |
| Muricidae | Ocenebrinae | *Eupleura muriciformis* |
| Muricidae | Ocenebrinae | *Mexacanthina angelica* |
| Muricidae | Ocenebrinae | *Mexacanthina lugubris* |
| Muricidae | Rapaninae | *Neorapana tuberculata* |
| Muricidae | Rapaninae | *Penethais triangularis* |
| Muricidae | Rapaninae | *Plicopurpura columellaris* |
| Muricidae | Ocenebrinae | *Roperia poulsoni* |
| Muricidae | Rapaninae | *Stramonita biserialis* |
| Muricidae | Ergalataxinae | *Trachypollia lugubris* |
| Muricidae | Rapaninae | *Vasula speciosa* |
| Muricidae | Aspellinae | *Zetecopsis zeteki* |
| Naticidae | Polinicinae | *Polinices uber* |
| Naticidae | Naticinae | *Notocochlis chemnitzii* |
| Neritidae | Neritinae | *Nerita funiculata* |
| Neritidae | Neritinae | *Nerita scabricosta* |
| Neritidae | Neritininae | *Vitta luteofasciata* |
| Olividae | Olivellinae | *Callianax biplicata* |
| Olividae | Olivellinae | *Olivella intorta* |
| Ovulidae | Cypraediinae | *Jenneria pustulata* |
| Phasianellidae |  | *Eulithidium sp 1.* |
| Pisaniidae |  | *Pusio elegans* |
| Rissoinidae |  | *Rissoina stricta* |
| Siphonariidae |  | *Siphonaria maura/palmata* |
| Tegulidae |  | *Agathistoma cortezianum* |
| Tegulidae |  | *Agathistoma eiseni* |
| Tegulidae |  | *Norrisia norrisii* |
| Tegulidae |  | *Tegula aureotincta* |
| Tegulidae |  | *Tegula funebralis* |
| Tegulidae |  | *Tegula gallina* |
| Tegulidae |  | *Tegula rugosa* |
| Terebridae | Terebrinae | *Neoterebra variegata* |
| Turbinidae | Turbininae | *Megaastrea undosa* |
| Turbinidae | Turbininae | *Pomaulax gibberosus* |
| Turbinidae | Turbininae | *Turbo fluctuosus* |
| Turbinidae | Turbininae | *Turbo funiculosus* |
| Turbinidae | Turbininae | *Uvanilla olivacea* |
| Vermetidae |  | Vermetidae *sp. 1* |
